# The autism-associated loss of δ-catenin functions disrupts social behaviors

**DOI:** 10.1101/2023.01.12.523372

**Authors:** Hadassah Mendez-Vazquez, Regan L. Roach, Kaila Nip, Matheus F. Sathler, Tyler Garver, Rosaline A. Danzman, Madeleine C. Moseley, Jessica P. Roberts, Olivia N. Koch, Ava A. Steger, Rahmi Lee, Jyothi Arikkath, Seonil Kim

**Author notes:** These authors contribute equally (Hadassah Mendez-Vazquez, Regan L. Roach, Kaila Nip). Department of Cellular and Integrative Physiology, University of Texas Health Science, Center San Antonio, San Antonio, TX, 78229. Corresponding Authors: Seonil Kim.

## Abstract

δ-catenin is expressed in excitatory synapses and functions as an anchor for the glutamatergic AMPA receptor (AMPAR) GluA2 subunit in the postsynaptic density. The glycine 34 to serine (G34S) mutation in the *δ-catenin* gene is found in autism spectrum disorder (ASD) patients and induces loss of δ-catenin functions at excitatory synapses, which is presumed to underlie ASD pathogenesis in humans. However, how the G34S mutation causes loss of δ-catenin functions to induce ASD remains unclear. Here, using neuroblastoma cells, we discover that the G34S mutation generates an additional phosphorylation site for glycogen synthase kinase 3β (GSK3β). This promotes δ-catenin degradation and causes the reduction of δ-catenin levels, which likely contributes to the loss of δ-catenin functions. Synaptic δ-catenin and GluA2 levels in the cortex are significantly decreased in mice harboring the δ-catenin G34S mutation. The G34S mutation increases glutamatergic activity in cortical excitatory neurons while it is decreased in inhibitory interneurons, indicating changes in cellular excitation and inhibition. δ-catenin G34S mutant mice also exhibit social dysfunction, a common feature of ASD. Most importantly, inhibition of GSK3β activity reverses the G34S-induced loss of δ-catenin function effects in cells and mice. Finally, using δ-catenin knockout mice, we confirm that δ-catenin is required for GSK3β inhibition-induced restoration of normal social behaviors in δ-catenin G34S mutant animals. Taken together, we reveal that the loss of δ-catenin functions arising from the ASD-associated G34S mutation induces social dysfunction via alterations in glutamatergic activity and that GSK3β inhibition can reverse δ-catenin G34S-induced synaptic and behavioral deficits.

**Significance Statement:** δ-catenin is important for the localization and function of glutamatergic AMPA receptors at synapses in many brain regions. The glycine 34 to serine (G34S) mutation in the *δ-catenin* gene is found in autism patients and results in the loss of δ-catenin functions. δ-catenin expression is also closely linked to other autism-risk genes involved in synaptic structure and function, further implying that it is important for the autism pathophysiology. Importantly, social dysfunction is a key characteristic of autism. Nonetheless, the links between δ-catenin functions and social behaviors are largely unknown. The significance of the current research is thus predicated on filling this gap by discovering the molecular, cellular, and synaptic underpinnings of the role of δ-catenin in social behaviors.

## Introduction

Social behaviors are essential for the survival of many species (1). However, our knowledge of the physiological, cellular, and molecular mechanisms driving social behaviors is still limited (1). Moreover, various mental disorders, including autism spectrum disorder (ASD), anxiety, depression, and schizophrenia, have social impairment as a primary symptom (2–5). Therefore, knowledge of the mechanisms mediating social behaviors can aid to better understand these diseases and develop effective treatments.

Postsynaptic glutamatergic activity in excitatory and inhibitory cells in the brain is crucial for social behaviors (6–10). In fact, disruptions in excitatory connectivity have been implicated in individuals with ASD and in mouse models of ASD (11–19). Therefore, maintaining the proper balance of cellular excitation and inhibition in the brain is essential to ensure normal social behaviors. However, it is unknown how postsynaptic glutamatergic activity in the brain regulates social behaviors. Several human genetic studies suggest that the *δ-catenin* gene is strongly linked to ASD (20–24). Notably, genetic alterations in the *δ-catenin* gene are associated with severely affected ASD patients from multiple families (21,23, 24). δ-catenin is a member of the armadillo repeat protein family that is highly expressed in neurons and localizes to the excitatory and inhibitory synapses (25–31). At the postsynaptic density (PSD) in the excitatory synapse, δ-catenin interacts with the intracellular domain of N-cadherin, a synaptic cell adhesion protein, and the carboxyl-terminus of δ-catenin binds to AMPA-type glutamate receptor (AMPAR)-binding protein (ABP) and glutamate receptor-interacting protein (GRIP) (**Fig. 1a**) (25, 28–30, 32, 33). This N-cadherin-δ-catenin-ABP/GRIP complex functions as an anchor for the AMPAR GluA2 subunit (**Fig. 1a**) and is disrupted when δ-catenin functions are lost (25, 30, 34, 35). Importantly, some δ-catenin ASD mutations are unable to reverse the δ-catenin deficiency-induced reduction of excitatory synapse density in cultured mouse neurons, suggesting that these mutations cause a loss of δ-catenin functions (24). A recent study reveals that δ-catenin knockdown increases the synaptic excitation and inhibition ratio and intrinsic excitability in young cortical pyramidal cells (31). Thus, this suggests that δ-catenin deficiency induces ASD-associated social deficits via impairment of glutamatergic synapses. Moreover, δ-catenin expression is closely linked to other ASD-risk genes involved in synaptic structure and function, such as GluA2, cadherins, GRIP, and synaptic Ras GTPase activating protein 1 (SYNGAP1), further implying that it is important for synaptic pathophysiology and social dysfunctions in ASD (31, 36–38). Nonetheless, the molecular and cellular links between δ-catenin functions, synaptic activity, and social behaviors are largely unknown.

**Figure 1.**
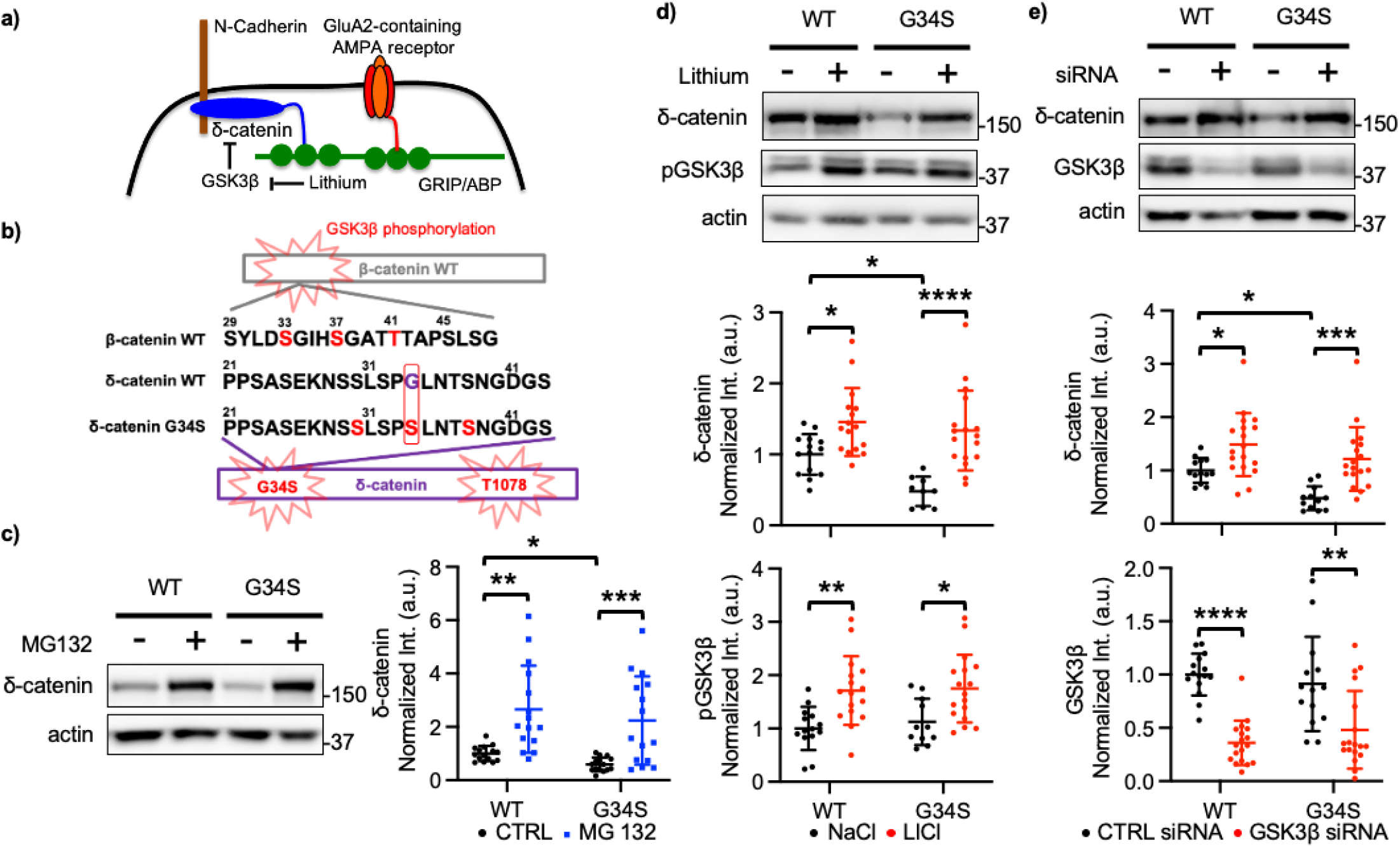
The δ-catenin G34S mutation increases GSK3β-mediated δ-catenin degradation. **a)** A schematic of N-cadherin-δ-catenin-ABP/GRIP-GluA2 synaptic complex and GSK3β regulation of δ-catenin. At PSD, δ-catenin interacts with N-cadherin, a synaptic cell adhesion protein. The carboxyl-terminus of δ-catenin binds to AMPA receptor-binding protein (ABP) and glutamate receptor-interacting protein (GRIP). This N-cadherin-δ-catenin-ABP/GRIP complex functions as an anchor for GluA2. GSK3β phosphorylates δ-catenin, which leads to δ-catenin degradation. The reduction of GSK3β activity by lithium stabilizes the N-cadherin-δ-catenin-ABP/GRIP-GluA2 complex. **b)** The δ-catenin G34S mutation adds an additional GSK3β-mediated phosphorylation site to induce δ-catenin degradation. GSK3β phosphorylation sites of β-catenin in the amino-terminal region known to induce proteasomal degradation of β-catenin highlighted in red (S33, S37, and T41) are comparable to possible GSK3β phosphorylation sites in the amino-terminus of G34S mutant δ-catenin also highlighted in red (S30, S34, and S38). In addition to GSK3β-mediated threonine 1078 phosphorylation site in the carboxyl-terminus of δ-catenin, these potential amino-terminal GSK3β-mediated phosphorylation sites may enhance δ-catenin degradation. **c)** Representative immunoblots and summary graphs of normalized δ-catenin levels in SH-SY5Y cell lysates transfected with WT or G34S δ-catenin in the presence (+) or absence (-) of 10 μM MG132 (n = 15 immunoblots from 5 independent cultures, Kruskal-Wallis test with the Dunn’s test, \**p* < 0.05, \*\**p* < 0.01, and \*\*\**p* < 0.001). **d)** Representative immunoblots and summary graphs of normalized δ-catenin and pGSK3β levels in SH-SY5Y cell lysates transfected with WT or G34S δ-catenin and treated with 2 mM NaCl (-) or 2 mM LiCl (+) (n = number of immunoblots from 5 independent cultures. For δ-catenin, WT + NaCl = 14, WT + LiCl = 16, G34S + NaCl = 9, and G34S + LiCl = 17. For pGSK3β, WT + NaCl = 14, WT + LiCl = 16, G34S + NaCl = 11, and G34S + LiCl = 17. Two-way ANOVA with the Tukey test, \**p* < 0.05, \*\**p* < 0.01, and \*\*\*\**p* < 0.0001). **e)** Representative immunoblots and summary graphs of normalized δ-catenin and GSK3β levels in SH-SY5Y cell lysates transfected with WT or G34S δ-catenin and treated with scrambled (CTRL) (-) or GSK3β (+) siRNA (n = number of immunoblots from 5 independent cultures. For δ-catenin, WT + CTRL siRNA = 13, WT + GSK3β siRNA = 18, G34S + CTRL siRNA = 12, and G34S + GSK3β siRNA = 18. For GSK3β, WT + CTRL siRNA = 15, WT + GSK3β siRNA = 18, G34S + CTRL siRNA = 14, and G34S + GSK3β siRNA = 18. Two-way ANOVA with the Tukey test, \**p* < 0.05, \*\**p* < 0.01, \*\*\**p* < 0.001, and \*\*\*\**p* < 0.0001). The position of molecular mass markers (kDa) is shown on the right of the blots.

Among the ASD-associated δ-catenin missense mutations, the glycine 34 to serine (G34S) mutation exhibits a profound loss of δ-catenin functions in excitatory synapses (24). However, how the δ-catenin G34S mutation causes the loss of δ-catenin functions to induce social deficits in ASD is not understood. Here, we reveal that the -catenin G34S mutation increases δ-catenin degradation by enhancing glycogen synthase kinase 3β (GSK3β)-mediated phosphorylation. Using δ-catenin G34S knockin mice, we find that the δ-catenin G34S mutation alters glutamatergic activity in cortical neurons and impairs social behaviors. Moreover, we show that a reduction of GSK3β activity can reverse the G34S-induced loss of δ-catenin function effects on δ-catenin levels, glutamatergic activity, and social behaviors. Finally, using δ-catenin knockout (KO) mice, we confirm that δ-catenin is required for GSK3β inhibition-induced restoration of normal social behavior in δ-catenin G34S animals. Importantly, recent studies suggest that changes in GSK3β activity may be an important aspect of the pathophysiology of ASD (39–46) Moreover, GSK3β is suggested to be a useful therapeutic target for brain disorders because GSK3β has been linked to cognitive processes (47). Therefore, our research not only discovers the molecular, cellular, and synaptic underpinnings of the role of δ-catenin in social behaviors but also identifies GSK3β as a new therapeutic target for limiting the loss of δ-catenin function-induced synaptopathy in ASD and related mental disorders.

## Results

### The δ-catenin G34S mutation increases GSK3β-mediated δ-catenin degradation

It has been previously shown that GSK3β phosphorylates threonine 1078 (T1078) of δ-catenin, leading to δ-catenin ubiquitination and subsequent proteasome-mediated degradation, whereas inhibition of GSK3β elevates δ-catenin levels (29, 48, 49) (**Fig. 1b**). GSK3β also phosphorylates β-catenin, another member of the armadillo-repeat proteins, and triggers its degradation in a similar manner (50, 51). In contrast to δ-catenin, which has a GSK3β-mediated phosphorylation site towards the carboxyl-terminus (T1078), β-catenin has the phosphorylation sites at serine 33 and 37 (S33 and S37) and threonine 41 (T41) in the amino-terminus (52) (**Fig. 1b**). The Group-based Prediction System (53) predicts that the G34S mutation in the amino-terminus of δ-catenin mimics one of the β-catenin phosphorylation sites, which may provide an additional site for GSK3β-mediated phosphorylation of δ-catenin (**Fig. 1b**). Thus, this may accelerate the degradation of δ-catenin, causing the loss of δ-catenin functions. To address whether the δ-catenin G34S mutation increases GSK3β-mediated δ-catenin degradation, we expressed wild-type (WT) or G34S δ-catenin in human neuroblastoma (SH-SY5Y) cells lacking endogenous δ-catenin expression and compared δ-catenin levels using immunoblotting. When normalized to WT δ-catenin, G34S δ-catenin levels were significantly reduced (WT, 1.000 ± 0.286 and G34S, 0.592 ± 0.260, *p* = 0.0227) (**Fig. 1c**). To determine whether reduced G34S δ-catenin levels were mediated by proteasomal degradation, we treated cells with 10 μM MG132, a proteasome inhibitor, for 4 hours and measured δ-catenin levels using immunoblotting. When proteasome activity was inhibited, both WT δ-catenin and mutant δ-catenin levels were significantly elevated in comparison to δ-catenin levels without MG132 treatment (WT + MG132, 2.659 ± 1.630, *p* = 0.0066, and G34S + MG132, 2.239 ± 1.658, *p* = 0.0002) (**Fig. 1c**). This suggests that the δ-catenin G34S mutation reduces δ-catenin levels, which is mediated by proteasomal degradation.

To examine whether reduced δ-catenin G34S expression was mediated by GSK3β, we used lithium chloride (LiCl) to inhibit GSK3β activity pharmacologically. We have previously shown that LiCl treatment significantly reduces GSK3β activity and increases δ-catenin expression in mouse cultured neurons and mouse brains (29). WT δ-catenin or G34S δ-catenin was expressed in SH-SY5Y cells, and these cells were treated with 2 mM LiCl or 2 mM sodium chloride (NaCl) for 18 hours before collecting whole cell lysates to analyze δ-catenin levels. We confirmed that mutant δ-catenin levels were significantly reduced in the absence of LiCl (WT + NaCl, 1.000 ± 0.287 and G34S + NaCl, 0.479 ± 0.208, *p* = 0.0354) (**Fig. 1d**). Importantly, LiCl treatment significantly elevated the levels of both WT δ-catenin and mutant δ-catenin (WT + LiCl, 1.456 ± 0.479, *p* = 0.0300, and G34S + LiCl, 1.336 ± 0.560, *p* < 0.0001) (**Fig. 1d**). Notably, we confirmed that LiCl treatment reduced GSK3β activity in cells by showing that it significantly increased phosphorylation at serine 9 of GSK3β (pGSK3β), an inactive form of GSK3β (48, 49), as seen previously (29) (WT + NaCl, 1.000 ± 0.405, WT + LiCl, 1.712 ± 0.644, *p* = 0.0051, G34S + NaCl, 1.124 ± 0.433, and G34S + LiCl, 1.749 ± 0.635, *p* = 0.0265) (**Fig. 1d**). To account for potential off-target effects from lithium treatment, we utilized siRNA for genetic inhibition of GSK3β. SH-SY5Y cells expressing WT δ-catenin or G34S δ-catenin were treated with either 25 nM scrambled siRNA as a control (CTRL) or 25 nM GSK3β siRNA for 72 hours. When we compared the WT δ-catenin and G34S δ-catenin levels in cells treated with CTRL siRNA, a significantly reduced δ-catenin level was observed in cells expressing mutant δ-catenin (WT + CTRL siRNA, 1.000 ± 0.231 and G34S + CTRL siRNA, 0.478 ± 0.223, *p* = 0.0429) (**Fig. 1e**). Both WT δ-catenin and G34S δ-catenin levels were significantly elevated by GSK3β siRNA-mediated knockdown (WT + GSK3β siRNA, 1.485 ± 0.592, *p* = 0.0365, and G34S + GSK3β siRNA, 1.215 ± 0.596, *p* = 0.0007) (**Fig. 1e**). We confirmed that GSK3β siRNA significantly reduced GSK3β levels, indicating a decrease in GSK3β activity (WT + CTRL siRNA, 1.000 ± 0.196, WT + GSK3β siRNA, 0.359 ± 0.208, *p* < 0.0001, G34S + CTRL siRNA, 0.912 ± 0.442, and G34S + GSK3β siRNA, 0.481 ± 0.365, *p* = 0.0017) (**Fig. 1e**). Taken together, we discover that the δ-catenin G34S mutation increases GSK3β-mediated δ-catenin degradation via a proteosome-mediated mechanism, likely leads to loss of δ-catenin functions. We further reveal that the reduction of GSK3β activity restores normal δ-catenin levels in cells expressing G34S δ-catenin.

### Additional GSK3β-mediated phosphorylation of G34S δ-catenin is important for increased δ-catenin degradation

Additional δ-catenin mutations were generated to test the hypothesis that the G34S mutation is phosphorylated by GSK3β to enhance δ-catenin degradation. First, we substituted the glycine 34 for alanine (A) to make a G34A mutant that was unable to be phosphorylated by GSK3β. We also generated a G34D mutant in which glycine was substituted with a phospho-mimetic aspartate (D) at position 34. WT δ-catenin, G34A δ-catenin, or G34D δ-catenin was expressed in SH-SY5Y cells, and δ-catenin levels were measured using immunoblotting. We found that there was no significant difference between WT δ-catenin and G34A δ-catenin expression, whereas G34D δ-catenin levels were significantly reduced, when compared with WT δ-catenin and G34A δ-catenin (WT, 1,000, G34A, 1.311 ± 0.437, and G34D, 0.662 ± 0.150, WT vs. G34A, *p* > 0.999, WT vs. G34D, *p* = 0.0088, G34A vs. G34D, *p* = 0.0088) (**Fig. 2a**), as seen in cells expressing G34S δ-catenin (**Fig. 1c-e**). To examine whether GSK3β-mediated phosphorylation of G34S δ-catenin is important for increased δ-catenin degradation, we first used LiCl to pharmacologically inhibit GSK3β activity as shown in **Fig. 1d**. Given that G34A δ-catenin acted similarly to WT δ-catenin (**Fig. 2a**), G34A δ-catenin or G34D δ-catenin was expressed in SH-SY5Y cells, and cells were treated with 2 mM LiCl or 2 mM NaCl for 18 hours. As seen in **Fig. 2a**, G34D δ-catenin levels were significantly reduced in the absence of LiCl when compared to G34A δ-catenin (G34A + NaCl, 1.000 ± 0.258 and G34D + NaCl, 0.551 ± 0.361, *p* = 0.0101) (**Fig. 2b**). LiCl treatment significantly elevated the levels of G34A δ-catenin, but not G34D δ-catenin (G34A + LiCl, 1.583 ± 0.458, *p* = 0.0005, and G34D + LiCl, 0.476 ± 0.255, *p* = 0.9433) (**Fig. 2b**). We confirmed that LiCl treatment significantly increased pGSK3β, an indication of GSK3β inhibition (G34A + NaCl, 1.000 ± 0.197, G34A + LiCl, 1.601 ± 0.531, *p* = 0.0317, G34D + NaCl, 0.982 ± 0.245, and G34D + LiCl, 1.557 ± 0.877, *p* = 0.0372) (**Fig. 2b**). Next, we used GSK3β siRNA for genetic inhibition of GSK3β activity. G34A δ-catenin or G34D δ-catenin was expressed in SH-SY5Y cells with either 25 nM scrambled siRNA (CTRL) or 25 nM GSK3β siRNA for 72 hours. When we compared the G34A δ-catenin and G34D δ-catenin levels in cells treated with scrambled siRNA, significantly reduced δ-catenin G34D levels were found (G34A + CTRL siRNA, 1.000 ± 0.166 and G34D + CTRL siRNA, 0.393 ± 0.230, *p* = 0.0330) (**Fig. 2c**). Like lithium treatment, GSK3β knockdown had no effect on G34D δ-catenin levels, but it significantly elevated G34A δ-catenin expression (G34A + GSK3β siRNA, 2.100 ± 0.841, *p* < 0.0001, and G34D + GSK3β siRNA, 0.594 ± 0.424, *p* = 0.8190) (**Fig. 2c**). As seen in **Fig. 1e**, GSK3β siRNA treatment significantly reduced GSK3β levels (G34A + CTRL siRNA, 1.000 ± 0.388, G34A + GSK3β siRNA, 0.467 ± 0.137, *p* = 0.0171, G34D + CTRL siRNA, 1.218 ± 0.599, and G34D + GSK3β siRNA, 0.479 ± 0.189, *p* = 0.0013) (**Fig. 2c**). This suggests the G34S mutation increases GSK3β-mediated phosphorylation of δ-catenin, thus enhances δ-catenin degradation, which likely contributes to loss of δ-catenin functions.

**Figure 2.**
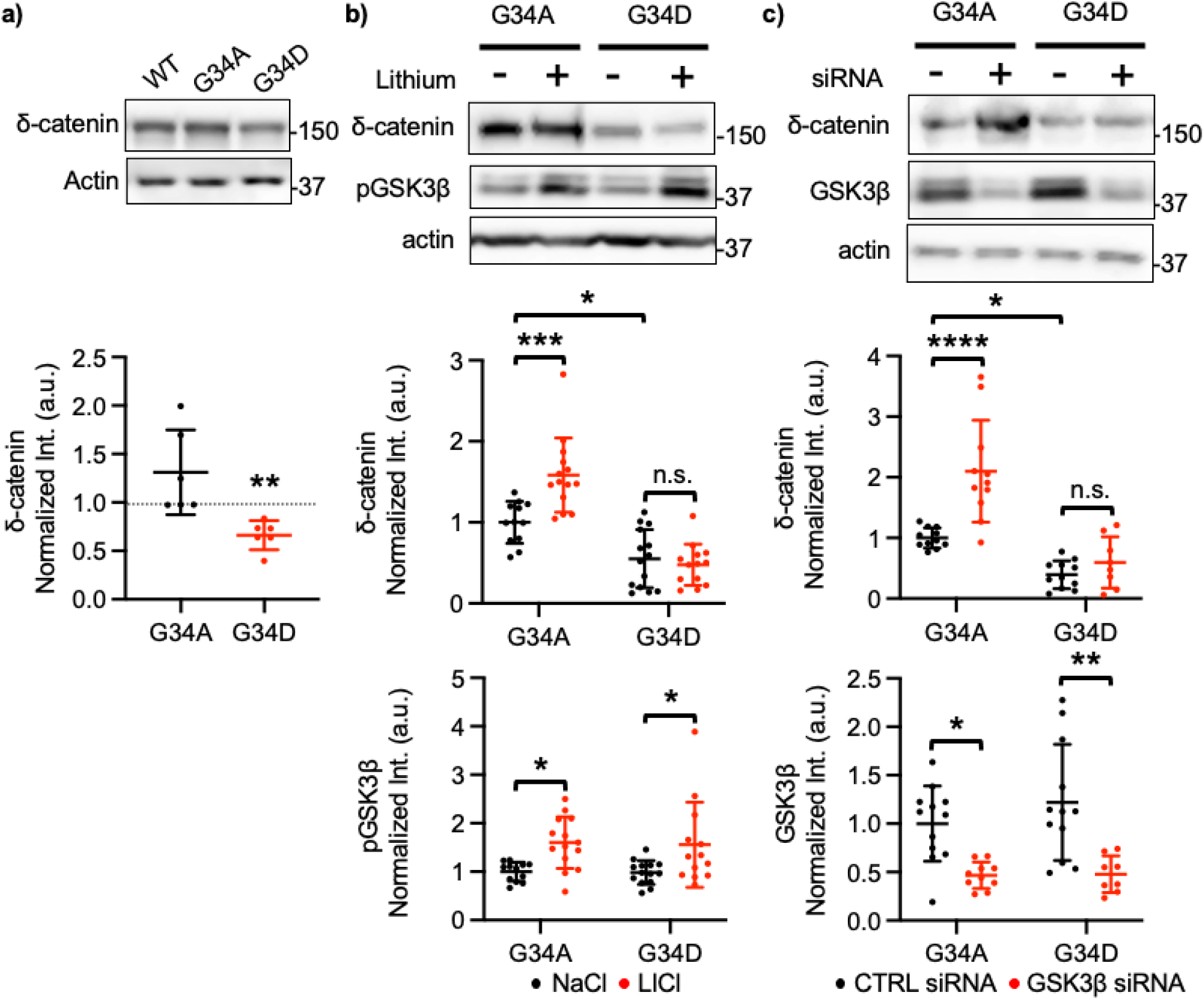
Additional GSK3β-mediated phosphorylation of G34S δ-catenin is important for increased δ-catenin degradation. **a)** Representative immunoblots and summary graphs of normalized δ-catenin levels in SH-SY5Y cell lysates transfected with WT, G34A, or G34D δ-catenin (n = 6 immunoblots from 3 independent cultures, Kruskal-Wallis test with the Dunn’s test, \*\**p* < 0.01). **b)** Representative immunoblots and summary graphs of normalized δ-catenin and pGSK3β levels in SH-SY5Y cell lysates transfected with G34A or G34D δ-catenin and treated with 2 mM NaCl (-) or 2 mM LiCl (+) (n = number of immunoblots from 3 independent cultures. G34A + NaCl = 12, G34A + LiCl = 14, G34D + NaCl = 14, and G34D + LiCl = 13. Two-way ANOVA with the Tukey test, \**p* < 0.05 and \*\*\**p* < 0.001. n.s. indicates no significant difference). **c)** Representative immunoblots and summary graphs of normalized δ-catenin and GSK3β levels in SH-SY5Y cell lysates transfected with WT or G34S δ-catenin and treated with scrambled (CTRL) (-) or GSK3β (+) siRNA (n = number of immunoblots from 3 independent cultures. For δ-catenin, G34A + CTRL siRNA = 11, G34A + GSK3β siRNA = 11, G34D + CTRL siRNA = 11, and G34D + GSK3β siRNA = 8. For GSK3β, G34A + CTRL siRNA = 12, G34A + GSK3β siRNA = 10, G34D + CTRL siRNA = 12, and G34D + GSK3β siRNA = 8. Two-way ANOVA with the Tukey test, \**p* < 0.05, \*\**p* < 0.01, and \*\*\*\**p* < 0.0001. n.s. indicates no significant difference). The position of molecular mass markers (kDa) is shown on the right of the blots.

### Lithium treatment reverses a significant reduction of synaptic δ-catenin and GluA2 in the δ-catenin G34S cortex

To understand how the G34S mutation affected synaptic δ-catenin and AMPAR levels in the brain, we used δ-catenin G34S knockin mice that were generated at University of Nebraska Medical Center (UNMC) Mouse Genome Engineering Core utilizing the CRISPR-Cas9 technique (54). The PSD fractions of the cortex were collected from 3-month-old female and male WT and δ-catenin G34S animals and analyzed for protein levels using immunoblotting. When compared to the WT cortex, we found a significant reduction in synaptic δ-catenin (WT female, 1.000, G34S female, 0.494 ± 0.334, *p* = 0.0070, WT male, 1.000, and G34S male, 0.494 ± 0.227, *p* = 0.0348) and GluA2 levels (WT female, 1.000, G34S female, 0.612 ± 0.192, *p* = 0.0017, WT male, 1.000, and G34S male, 0.540 ± 0.222, *p* = 0.0440) in the female and male G34S animals’ cortices, but no change in synaptic GluA1 levels (WT female, 1.000, G34S female, 0.943 ± 0.619, *p* = 0.4723, WT male, 1.000, and G34S male, 0.848 ± 0.206, *p* = 0.3362) (**Fig. 3a and 3b**). Next, synaptic δ-catenin and AMPAR levels in the hippocampus were similarly analyzed using immunoblotting. We found no difference in synaptic δ-catenin (WT female, 1.000, G34S female, 0.929 ± 0.222, *p* = 0.4344, WT male, 1.000, and G34S male, 0.847 ± 0.371, *p* = 0.4843), GluA1 (WT female, 1.000, G34S female, 1.293 ± 0.457, *p* = 0.3876, WT male, 1.000, and G34S male, 0.880 ± 0.311, *p* = 0.8050), and GluA2 levels (WT female, 1.000, G34S female, 0.931 ± 0.366, *p* = 0.3654, WT male, 1.000, and G34S male, 0.848 ± 0.206, *p* = 0.9672) between the WT and G34S hippocampus (**Fig. S1a**). This suggests that the δ-catenin G34S mutation significantly reduces synaptic δ-catenin and GluA2 levels selectively in the cortex, but not in the hippocampus. This difference is likely due to higher GluA2 levels in the mouse hippocampus than the cortex (55, 56), thus AMPAR levels in the hippocampus is less affected by the δ-catenin G34S mutation compared to cortical AMPAR levels.

**Figure 3.**
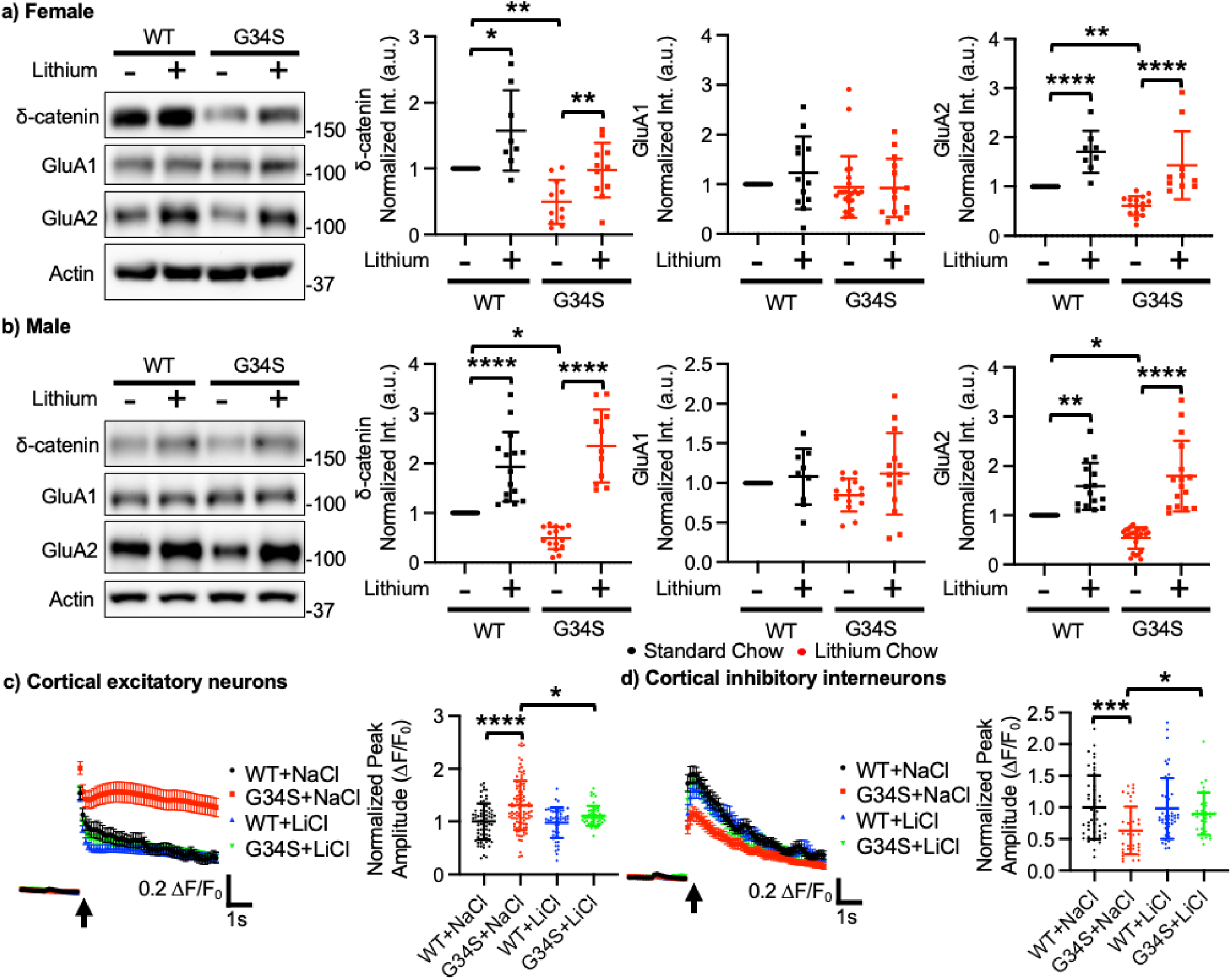
Lithium treatment reverses a significant reduction of synaptic δ-catenin and GluA2 in the δ-catenin G34S cortex and alterations in glutamatergic activity in cultured cortical neurons. Representative immunoblots and summary graphs of normalized δ-catenin, GluA1, and GluA2 levels in the cortical PSD fractions from standard chow (-) or lithium chow (+)-fed WT and δ-catenin G34S **a)** female (n = number of immunoblots [number of mice]. For δ-catenin, WT + standard chow =14 [5], WT + lithium chow = 8 [5], G34S + standard chow = 11 [6], and G34S + lithium chow = 11 [6]. For GluA1, WT + standard chow = 14 [5], WT + lithium chow = 12 [5], G34S + standard chow = 23 [6], and G34S + lithium chow = 14 [6]. For GluA2, WT + standard chow = 11 [5], WT + lithium chow = 8 [5], G34S + standard chow = 16 [6], and G34S + lithium chow = 10 [6]) and **b)** male mice (n = number of immunoblots [number of mice]. For δ-catenin, WT + standard chow = 17 [5], WT + lithium chow = 15 [5], G34S + standard chow = 15 [6], and G34S + lithium chow = 11 [6]. For GluA1, WT + standard chow = 14 [5], WT + lithium chow = 9 [5], G34S + standard chow = 14 [6], and G34S + lithium chow = 14 [6]. For GluA2, WT + standard chow = 18 [5], WT + lithium chow = 15 [5], G34S + standard chow = 24 [6], and G34S + lithium chow = 15 [6], \**p* < 0.05, \*\**p* < 0.01, and \*\*\*\**p* < 0.0001, Two-way ANOVA with the Tukey test). The position of molecular mass markers (kDa) is shown on the right of the blots. **c)** Average traces of GCaMP7s signals and summary data of normalized peak amplitude in each condition in excitatory neurons (n = number of neurons from 3 independent cultures, WT + NaCl = 75, G34S + NaCl = 98, WT + LiCl = 50, and G34S + LiCl = 49). **d)** Average traces of GCaMP6f signals and summary data of normalized peak amplitude in each condition in inhibitory interneurons (n = number of neurons from 3 independent cultures, WT + NaCl = 46, G34S + NaCl = 40, WT + LiCl = 46, and G34S + LiCl = 31). \**p* < 0.05, \*\*\**p* < 0.01, and \*\*\*\**p* < 0.0001, Two-way ANOVA with the Tukey test). An arrow indicates photostimulation.

We next examined if lithium-based pharmacological inhibition of GSK3β activity *in vivo* reversed the human autism-associated δ-catenin G34S mutation’s effects on synaptic δ-catenin and AMPAR levels in the cortex. For lithium treatment, we fed 3-month-old female and male WT and δ-catenin G34S mice with standard chow containing 0.17% w/w lithium carbonate *ad libitum* for 10 days, a protocol previously shown to significantly inhibit GSK3β activity and elevate δ-catenin levels in the brain (29). After treatment, whole cell lysates were collected from the cortex, and immunoblots of pGSK3β were used to assess whether GSK3β activity was inhibited in the cortex. As expected, in the male and female δ-catenin G34S cortex, lithium treatment significantly increased pGSK3β, when compared to the controls (WT female + standard chow, 1.000 ± 0.311, WT female + lithium chow, 1.495 ± 0.292, *p* = 0.0175, G34S female + standard chow, 1.024 ± 0.200, G34S female + lithium chow, 1.481 ± 0.551, *p* = 0.0222, WT male + standard chow, 1.000 ± 0.118, WT male + lithium chow, 1.455 ± 0.355, *p* = 0.0087, G34S male + standard chow, 0.931 ± 0.347, and G34S male + lithium chow, 1.347 ± 0.314, *p* = 0.0006), confirming the reduction of GSK3β activity *in vivo* (**Fig. S2a and S2b**). Next, the PSD fractions of the cortex were collected from 3-month-old female and male WT and δ-catenin G34S animals after lithium treatment to determine if the reduction of GSK3β activity restored normal synaptic δ-catenin and GluA2 levels using immunoblotting as seen in neuroblastoma cells (**Fig. 1d**). We found a reduction of synaptic δ-catenin (WT female + lithium chow, 1.578 ± 0.611, *p* = 0.0189, G34S female + lithium chow, 0.977 ± 0.414, *p* = 0.0088, WT male + lithium chow, 1.929 ± 0.699, *p* < 0.0001, and G34S male + lithium chow, 2.350 ± 0.734, *p* < 0.0001) and GluA2 levels (WT female + lithium chow, 1.705 ± 0.431, *p* < 0.0001, G34S female + lithium chow, 1.432 ± 0.693, *p* < 0.0001, WT male + lithium chow, 1.588 ± 0.476, *p* = 0.0016, and G34S male + lithium chow, 1.795 ± 0.714, *p* < 0.0001) in the mutant female and male cortex was reversed by lithium treatment (**Fig. 3a and 3b**). However, we found no difference in GluA1 levels (WT female + lithium chow, 1.232 ± 0.731, *p* = 0.7081, G34S female + lithium chow, 0.926 ± 0.587, *p* = 0.8723, WT male + lithium chow, 1.078 ± 0.355, *p* = 0.7834, and G34S male + lithium chow, 1.116 ± 0.515, *p* = 0.0539) between the WT and G34S cortex following lithium treatment (**Fig. 3a and 3b**). Our findings indicate that the pharmacological inhibition of GSK3β activity is sufficient to restore normal synaptic δ-catenin and GluA2 levels in the G34S mutant cortex.

### Altered glutamatergic activity in cultured δ-catenin G34S cortical neurons

Certain types of ASD may be caused by a decrease in the signal-to-noise ratio in important brain circuits, which may be brought on by changes in a neuronal excitation and inhibition (E/I) balance (57). Moreover, postsynaptic glutamatergic activity in cortical excitatory or inhibitory neurons controls social behavior via the regulation of neuronal E/I within the cortical neural network (13, 19). Consistently, a recent study shows that δ-catenin deficiency increases the E/I ratio and intrinsic excitability in young cortical excitatory cells (31). Given that GluA2 levels are significantly reduced in the δ-catenin G34S cortex (**Fig. 3a and 3b**), we carried out Ca^2+^ imaging with glutamate uncaging in cultured primary WT and δ-catenin G34S mouse cortical neurons to determine if δ-catenin G34S changed glutamatergic activity in excitatory and inhibitory cortical cells. Glutamatergic activity was significantly increased in cultured 12-14 days *in vitro* (DIV) mouse excitatory cortical δ-catenin G34S neurons compared to WT cells (WT, 1.000 ± 0.340 ΔF/F_0_ and G34S, 1.307 ± 0.465 ΔF/F_0_, *p* < 0.0001) (**Fig. 3c**). Conversely, glutamatergic activity in cultured G34S inhibitory interneurons was markedly lower compared to WT cells (WT, 1.000 ± 0.504 ΔF/F_0_ and G34S, 0.621 ± 0.360 ΔF/F_0_, *p* = 0.0005) (**Fig. 3d**), suggesting that glutamatergic activity in δ-catenin G34S excitatory and inhibitory cells is altered in opposite ways. As the pharmacological inhibition of GSK3β activity restore normal synaptic expression of δ-catenin and GluA2 in the cortex (**Fig. 3a and 3b**), we further determined whether lithium treatment was able to reverse changes in glutamatergic activity in mutant neurons. We treated cultured neurons with 2 mM LiCl for 4 hours then performed Ca^2+^ imaging with glutamate uncaging. Pharmacological inhibition of GSK3β activity reversed the G34S δ-catenin effects on glutamatergic activity in cultured cortical excitatory (G34S, 1.109 ± 0.188 ΔF/F_0_, *p* = 0.0109) and inhibitory cells (G34S, 0.898 ± 0.334 ΔF/F_0_, *p* < 0.0419) (**Fig. 3c and 3d**). However, lithium treatment in WT neurons had no effect on glutamatergic activity (Excitatory neurons, WT + LiCl, 0.978 ± 0.290 ΔF/F_0_, *p* = 0.9869, and inhibitory interneurons, WT + LiCl, 0.982 ± 0.481 ΔF/F_0_, *p* = 0.9973) (**Fig. 3c and 3d**). Taken together, we demonstrate that δ-catenin G34S significantly alters glutamatergic activity in excitatory and inhibitory cells in opposite ways, which likely disrupts the neuronal E/I balance in the cortex. Moreover, we find that these changes in G34S mutant neurons are reversed by pharmacological inhibition of GSK3β.

### δ-catenin G34S induces social dysfunction in mice, which is reversed by GSK3β inhibition

A three-chamber test was performed to determine whether δ-catenin G34S affected social behaviors. We discovered that WT female and male mice interacted significantly longer with stranger 1 than the novel object during the sociability test, an indication of normal sociability (**Fig. 4a and Table S1**). Conversely, in δ-catenin G34S female and male mice, no significant difference in total interaction time was found between stranger 1 and the novel object (**Fig. 4a and Table S1**). Importantly, social interaction time with stranger 1 in δ-catenin G34S female and male mice was significantly less than WT mice (**Fig. 4a and Table S1**). In the social novelty test, WT mice engaged with stranger 2 for longer than they did with stranger 1, demonstrating normal social novelty (**Fig. 4b and Table S1**). However, there was no significant difference in reciprocal sniffing time between stranger 1 and stranger 2 in δ-catenin G34S female and male mice (**Fig. 4b and Table S1**). Crucially, compared to WT mice, social interaction with stranger 2 was markedly lower in both female and male δ-catenin G34S animals (**Fig. 4b and Table S1**). These findings indicate that δ-catenin G34S mice exhibit disrupted sociability and social novelty in the three-chamber test.

**Figure 4.**
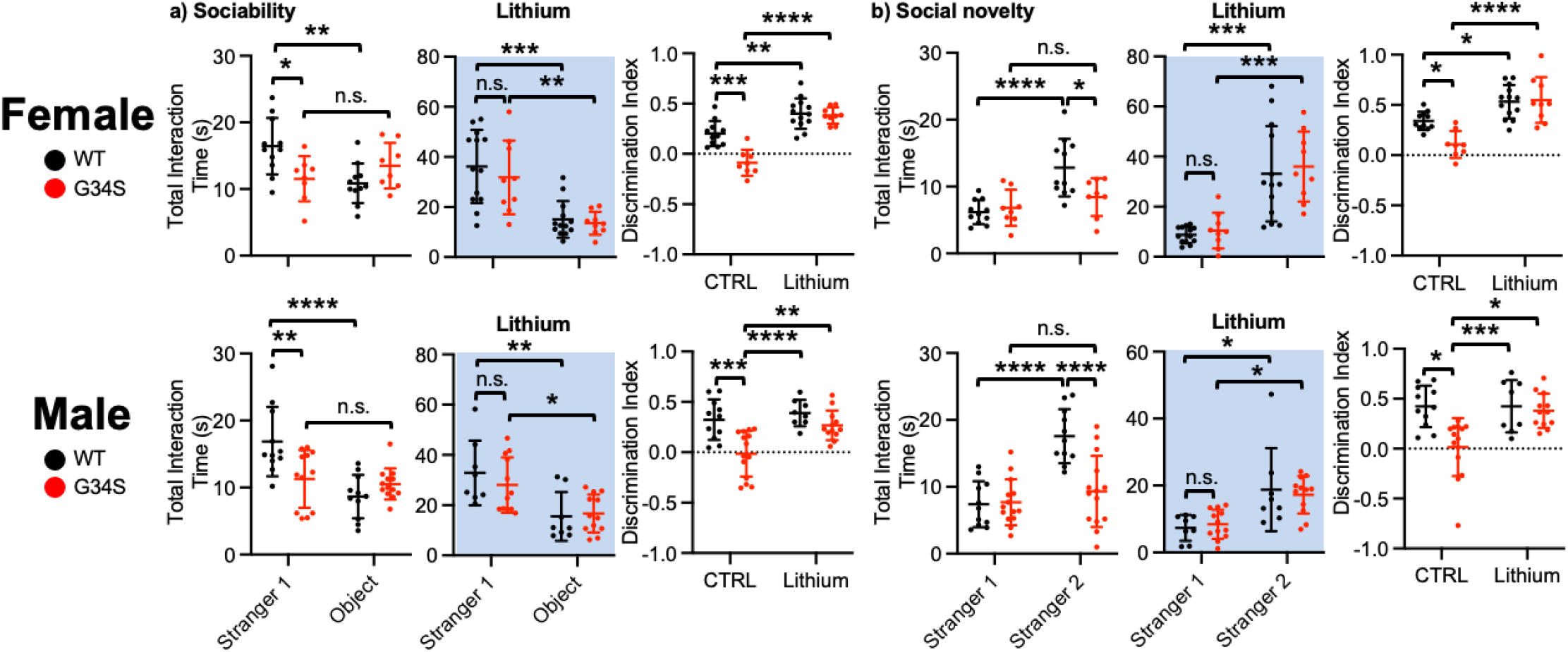
δ-catenin G34S induces social dysfunction in mice, which is reversed by GSK3β inhibition. Total interaction time and the discrimination index of **a)** sociability and **b)** social novelty in female and male mice in each genotype in control and lithium treated conditions (highlighted in light blue) in the three-chamber test (n = number of animals, WT female + standard chow = 11, WT female + lithium chow = 9, G34S female + standard chow = 8, G34S female + lithium chow = 7, WT male + standard chow =11, WT male + lithium chow = 8, G34S male + standard chow = 14, and G34S male + lithium chow = 12, \**p* < 0.05, \*\**p* < 0.01, \*\*\**p* < 0.001, and \*\*\*\**p* < 0.0001, Two-way ANOVA with Tukey test. n.s. indicates no significant difference).

As lithium treatment significantly reduced GSK3β activity and elevated synaptic δ-catenin and GluA2 expression in the δ-catenin G34S cortex (**Fig. 3a and 3b and Fig. S2**), we examined whether lithium treatment could restore normal social behaviors in mutant mice using the three-chamber test. In the sociability test, we discovered that both WT and δ-catenin G34S female and male mice fed with lithium chow interacted with stranger 1 for significantly longer than they did with the novel object, indicating that reduced GSK3β activity reversed disrupted sociability in δ-catenin G34S mice (**Fig. 4a and Table S1**). We further found that decreased social interaction time with stranger 1 in δ-catenin G34S mice was significantly reversed by lithium treatment in the sociability test (**Fig. 4a and Table S1**). The discrimination index further confirmed that lithium treatment reversed impaired sociability in δ-catenin G34S female and male mice (**Fig. 4a and Table S2**). We also revealed that lithium-fed WT and δ-catenin G34S female and male mice interacted significantly longer with stranger 2 than stranger 1 during the social novelty test, showing that lithium treatment restored normal social novelty preference in δ-catenin G34S mice (**Fig. 4b and Table S1**). In addition, reduced social interaction with stranger 2 in δ-catenin G34S mice was significantly elevated by lithium treatment in the social novelty test (**Fig. 4b and Table S1**). Finally, the discrimination index confirmed that lithium treatment restored normal social novelty behavior in δ-catenin G34S female and male mice (**Fig. 4b and Table S2**). Taken all together, δ-catenin G34S disrupts social behaviors in mice, which is reversed by lithium-based pharmacological inhibition of *in vivo* GSK3β activity.

### δ-catenin G34S mice show normal olfaction, anxiety levels, and locomotor activity

It is possible that sensory deficits may affect performances in social behavioral assays (58). During social activity, mice substantially rely on olfaction (58). The buried food test is a reliable method that depends on the mouse’s natural desire to forage using olfactory cues (59). Thus, we conducted the buried food test to examine whether the δ-catenin G34S mutation affected olfactory functions. We found no significant difference in the latency to find the buried food in δ-catenin G34S female and male mice compared to WT littermates (**Fig. S3a and Table S3**), suggesting that δ-catenin G34S mice have normal olfaction. Anxiety levels can also affect social behaviors in animals (60, 61). We thus measured locomotor activity and anxiety-like behavior using the open field test. We found no difference between genotype groups regardless of sex in total traveled distance (locomotor activity) and time spent outside and inside (anxiety-like behavior) (**Fig. S3b and Table S3**). This indicates that δ-catenin G34S mice have normal locomotor activity and anxiety levels, thus altered behaviors observed in the three-chamber test (**Fig. 4a and 4b**) strongly indicates social dysfunction.

### A significant reduction of synaptic GluA2 in the δ-catenin KO cortex and altered glutamatergic activity in cultured δ-catenin KO cortical neurons

We additionally generated δ-catenin KO mice to further confirm whether the loss of δ-catenin functions was important for glutamatergic activity in cortical neurons and social behaviors in animals. We examined if δ-catenin KO affected synaptic AMPAR levels in the brain. First, we confirmed via immunoblotting that there was no δ-catenin expression found in the cortex and hippocampus of δ-catenin KO female and male mice (**Fig. 5a and 5b and Fig. S1b**). The PSD fractions of the cortex were collected from 3-month-old female and male WT and δ-catenin KO animals and carried out immunoblotting to analyze for synaptic AMPAR levels as shown in **Fig. 3a and 3b**. When compared to the WT cortex, we found no difference in synaptic GluA1 levels (WT female, 1.000, KO female, 1.074 ± 0.268, *p* = 0.9426, WT male, 1.000, and KO male, 0.996 ± 0.089, *p* = 0.8290) but a significant reduction in GluA2 levels (WT female, 1.000, KO female, 0.757 ± 0.206, *p* = 0.0256, WT male, 1.000, and KO male, 0.754 ± 0.187, *p* = 0.0175) in the female and male KO animals’ cortices (**Fig. 5a and 5b**), as seen in the G34S mouse cortex (**Fig. 3a and 3b**). Next, the PSD fractions of the hippocampus were collected from 3-month-old female and male WT and δ-catenin KO mice and analyzed for synaptic AMPAR levels using immunoblotting. Like the δ-catenin G34S hippocampus, we found no difference in synaptic GluA1 (WT female, 1.000, KO female, 0.895 ± 0.092, *p* = 0.1870, WT male, 1.000, and KO male, 0.866 ± 0.086, *p* = 0.8050) and GluA2 levels (WT female, 1.000, KO female, 0.9929 ± 0.119, *p* = 0.1134, WT male, 1.000, and KO male, 0.882 ± 0.103, *p* = 0.1860) between the WT and KO hippocampus (**Fig. S1b**). This suggests that the loss of δ-catenin expression in mice significantly reduces synaptic GluA2 levels selectively in the cortex, but not in the hippocampus, which likely alters AMPAR-mediated glutamatergic activity in the δ-catenin KO cortex, like the G34S cortex. Thus, we conducted Ca^2+^ imaging with glutamate uncaging in cultured WT and δ-catenin KO mouse cortical neurons to determine if δ-catenin KO altered glutamatergic activity in excitatory and inhibitory cortical neurons as shown in **Fig 3c and 3d**. Glutamate-induced Ca^2+^ activity was markedly elevated in cultured 12-14 DIV mouse excitatory cortical δ-catenin KO neurons compared to WT cells (WT, 1.000 ± 0.274 ΔF/F_0_ and KO, 1.275 ± 0.230 ΔF/F_0_, *p* < 0.0001) (**Fig. 5c**). Conversely, glutamatergic activity in cultured KO inhibitory interneurons was significantly lower than WT cells (WT, 1.000 ± 0.0.472 ΔF/F_0_ and KO, 0.635 ± 0.327 ΔF/F_0_, *p* = 0.0005) (**Fig. 5d**). This suggests that glutamatergic activity in δ-catenin KO excitatory and inhibitory cells is altered in opposite ways, as seen in G34S cortical neurons (**Fig. 3c and 3d)**.

**Figure 5.**
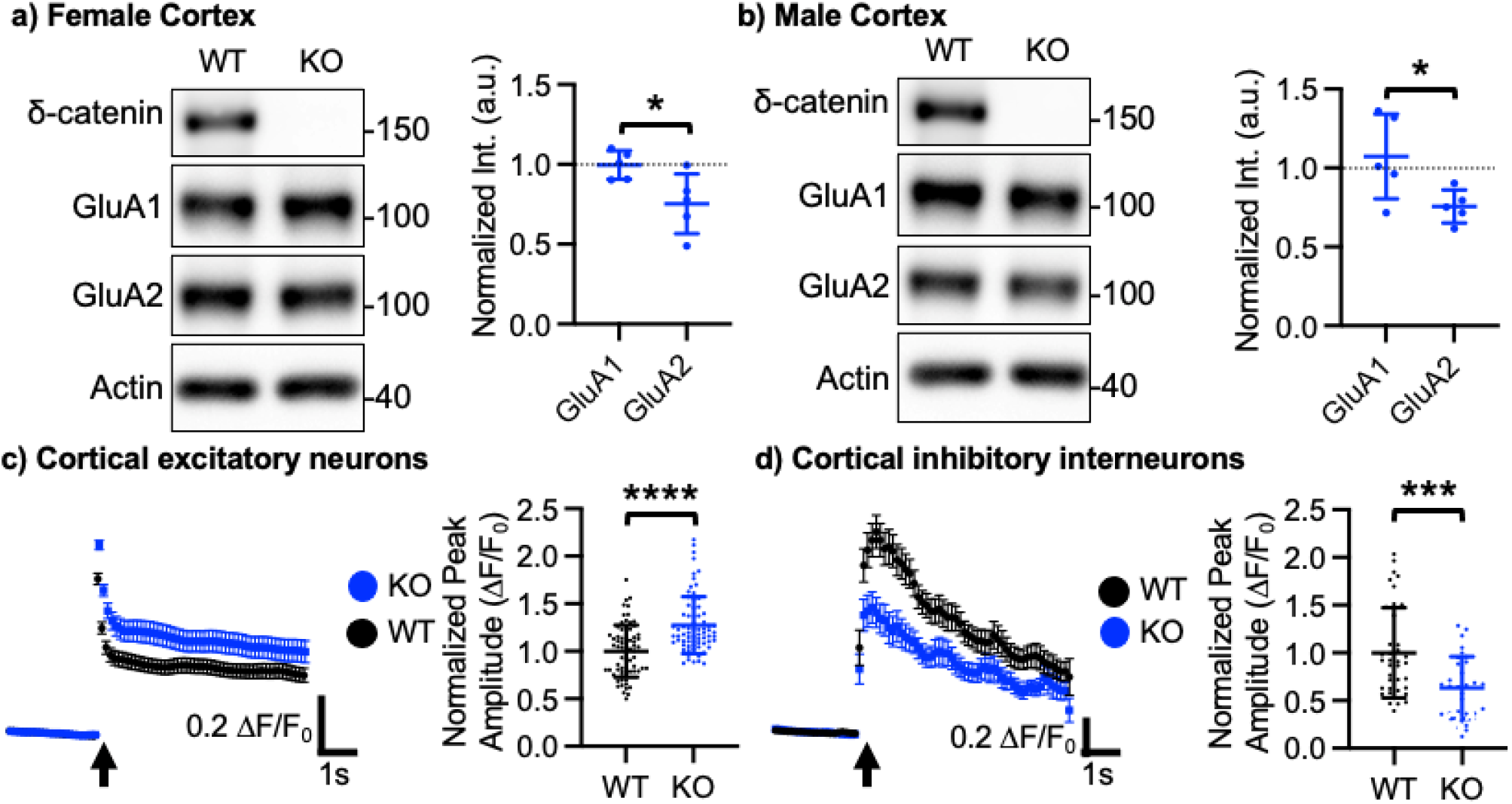
A significant reduction of synaptic GluA2 in the δ-catenin KO cortex and altered glutamatergic activity in cultured δ-catenin KO cortical neurons. Representative immunoblots and summary graphs of normalized GluA1, and GluA2 levels in the cortical PSD fractions from WT and δ-catenin KO **a)** female and **b)** male mice (n = 5 immunoblots from 3 mice in each condition, Kruskal-Wallis test with the Dunn’s test, \**p* < 0.05). No δ-catenin expression was found in the KO cortex. The position of molecular mass markers (kDa) is shown on the right of the blots. **c)** Average traces of GCaMP7s signals and summary data of normalized peak amplitude in each condition in in excitatory neurons (n = number of neurons from 3 independent cultures, WT = 70 and KO = 73). **d)** Average traces of GCaMP6f signals and summary data of normalized peak amplitude in each condition in inhibitory interneurons (n = number of neurons from 3 independent cultures, WT = 38 and KO = 30). \*\*\**p* < 0.01 and \*\*\*\**p* < 0.0001, the unpaired two-tailed Student’s t-test). An arrow indicates photostimulation

### Abnormal social behavior but normal olfaction, locomotor activity, and anxiety levels in δ-catenin KO mice

We used the three-chamber test to determine whether the lack of δ-catenin expression affected social behaviors in mice. In the sociability test, WT female and male mice interacted significantly longer with stranger 1 than a novel object, showing normal sociability (**Fig. 6a and Table S4**). However, in δ-catenin KO female and male mice, no significant difference in total interaction time was found between stranger 1 and the novel object (**Fig. 6a and Table S4**). Importantly, social interaction with stranger 1 in δ-catenin KO female and male mice was markedly lower than WT mice (**Fig. 6a and Table S4**). In the social novelty test, WT mice interacted with stranger 2 for longer than they did with stranger 1, an indication of normal social novelty preference (**Fig. 6b and Table S4**). Conversely, there was no significant difference in reciprocal sniffing time between stranger 1 and stranger 2 in δ-catenin KO female and male mice (**Fig. 6b and Table S4**). Compared to WT animals, social interaction with stranger 2 was significantly lower in both female and male δ-catenin KO mice (**Fig. 6b and Table S4**). Moreover, the discrimination index further confirmed that δ-catenin KO disrupted social behaviors in female and male mice (**Fig. 6a and 6b and Table S5**). These findings indicate that δ-catenin KO mice exhibit abnormal sociability and social novelty in the three-chamber test like δ-catenin G34S mice. In addition, we found that δ-catenin KO female and male mice showed normal olfactory function in the buried food test (**Fig. S3c and Table S3**). Finally, the open field test showed normal locomotor activity and anxiety levels in δ-catenin KO female and male mice (**Fig. S3d and Table S3**). This indicates that δ-catenin KO mice have normal locomotor activity and anxiety levels. Altogether, we conclude that δ-catenin KO mice exhibit abnormal social behaviors in the three-chamber test without altering olfaction, locomotor activity, and anxietylike behavior.

**Figure 6.**
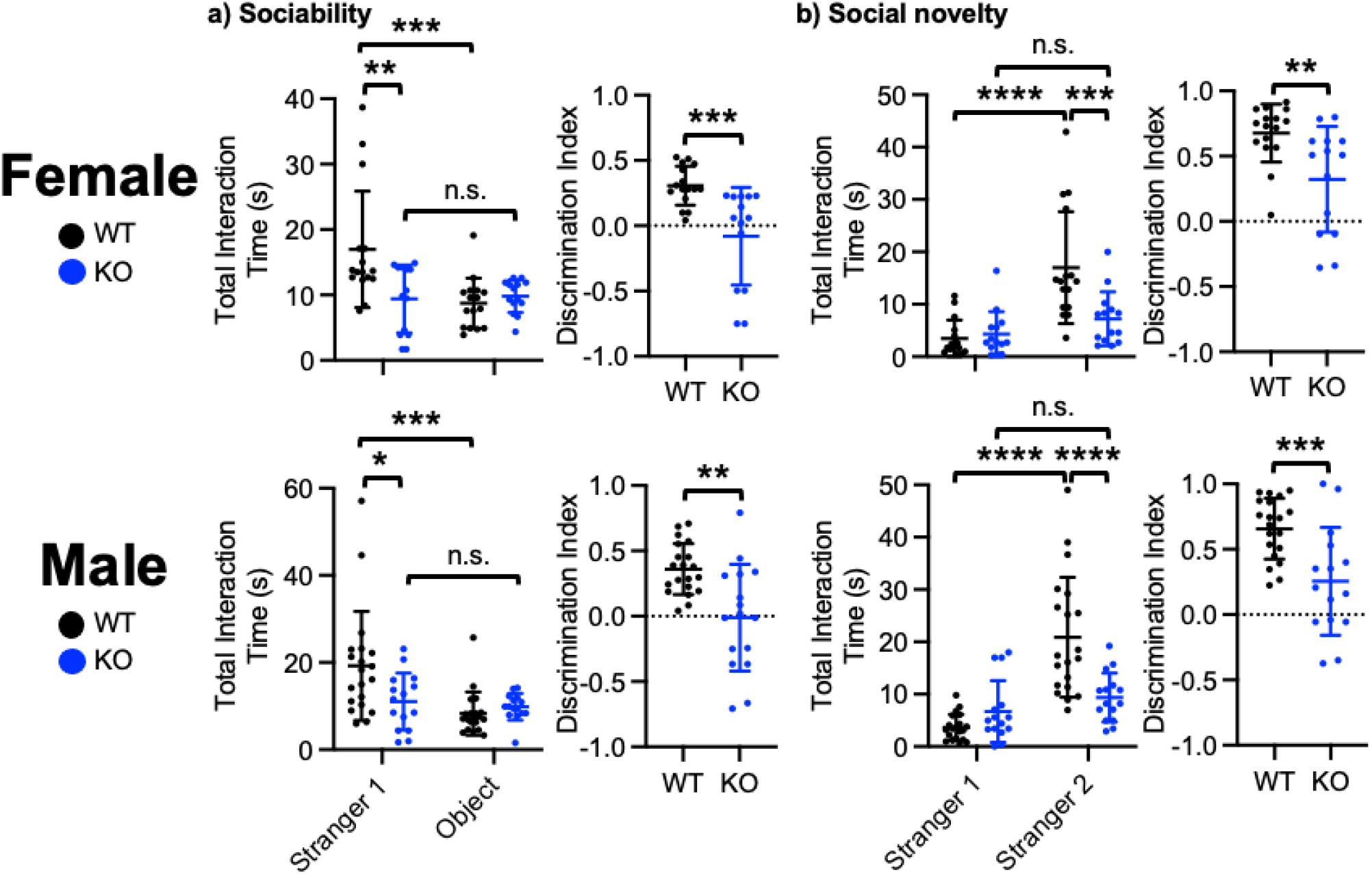
δ-catenin KO induces social dysfunction in mice. Total interaction time and the discrimination index of **a)** sociability and **b)** social novelty in female and male WT and KO mice in the three-chamber test (n = number of animals, WT female = 16, KO female = 14, WT male = 20, and KO male = 15, \**p* < 0.05, \*\**p* < 0.01, \*\*\**p* < 0.001, and \*\*\*\**p* < 0.0001, For total interaction time, Twoway ANOVA with Tukey test. For the discrimination index, the unpaired two-tailed Student’s t-test. n.s. indicates no significant difference).

### δ-catenin is required for lithium-induced restoration of normal social behavior in G34S mice

Lastly, we used δ-catenin KO mice to address whether δ-catenin was required for lithium effects on social behaviors. We fed δ-catenin KO mice with lithium chow and conducted the three-chamber test and compared their social behaviors to that of lithium-treated δ-catenin G34S animals. In the sociability test, we found no significant difference in total interaction time between stranger 1 and the novel object in lithium-treated δ-catenin KO female and male mice, whereas we found normal sociability in lithium-fed δ-catenin G34S animals (**Fig. 7a and Table S6**). Importantly, social interaction with stranger 1 in δ-catenin KO female and male mice was markedly lower than G34S mice following lithium treatment (**Fig. 7a and Table S6**). In the social novelty test, after lithium was administered, there was no significant difference in reciprocal sniffing time between stranger 1 and stranger 2 in δ-catenin KO female and male mice, while G34S female animals exhibited normal social novelty following lithium treatment (**Fig. 7b and Table S6**). Finally, the discrimination index revealed that lithium was unable to restore normal sociability and social novelty in δ-catenin KO female and male mice unlike δ-catenin G34S animals (**Fig. 7a and 7b and Table S7**). These findings suggest that δ-catenin is required for lithium-induced restoration of normal social behaviors in δ-catenin G34S mice.

**Figure 7.**
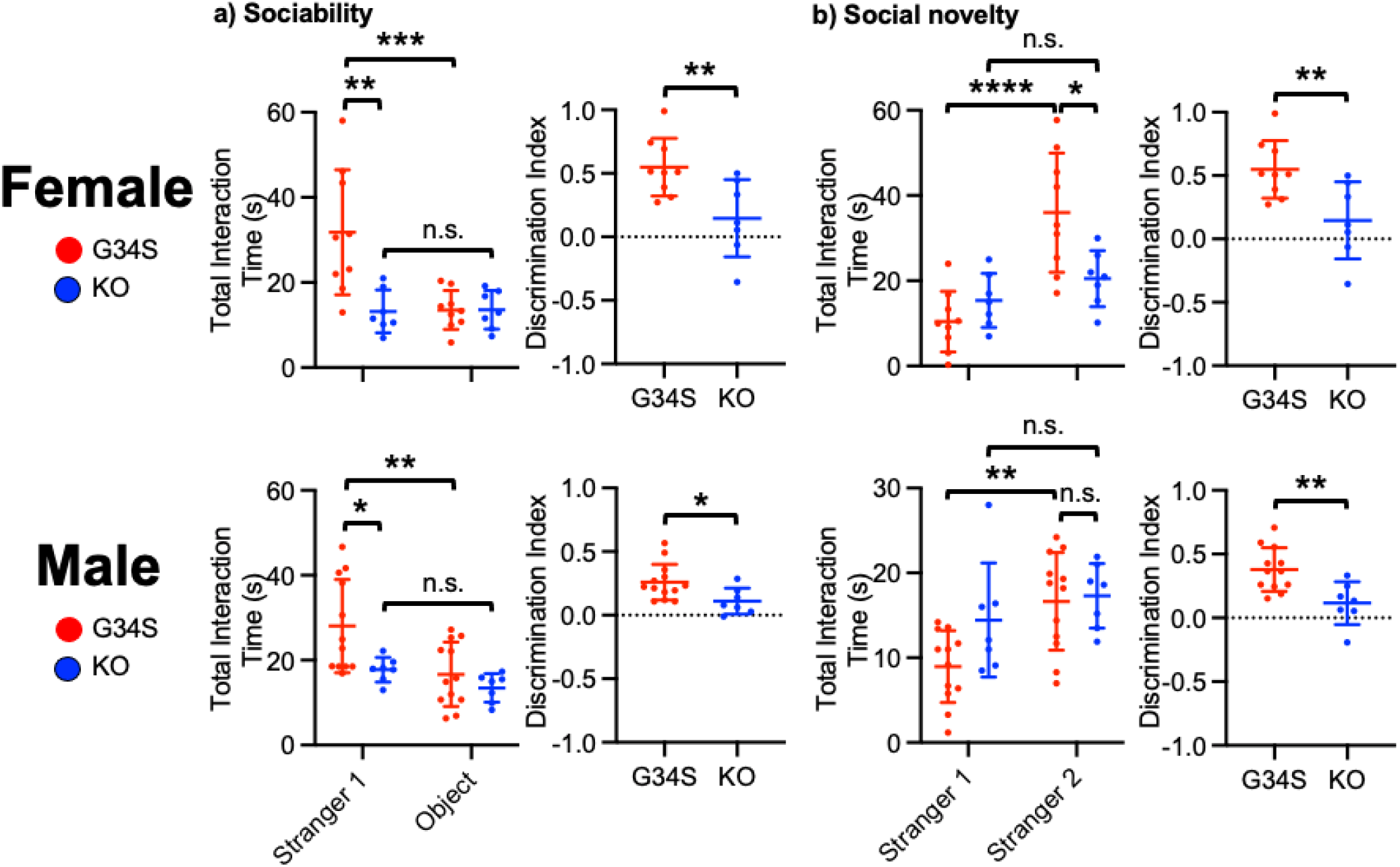
δ-catenin is required for lithium-induced restoration of normal social behaviors in G34S mice. Total interaction time and the discrimination index of **a)** sociability and **b)** social novelty in lithium-treated female and male G34S and KO mice in the three-chamber test (n = number of animals, G34S female + lithium = 9, KO female + lithium = 7, G34S male + lithium = 12, and KO male + lithium = 7, \**p* < 0.05, \*\**p* < 0.01, \*\*\**p* < 0.001, and \*\*\*\**p* < 0.0001, For total interaction time, Two-way ANOVA with Tukey test. For the discrimination index, the unpaired two-tailed Student’s t-test. n.s. indicates no significant difference).

## Discussion

ASD is a multifactorial neurodevelopmental disorder characterized by impaired social communication, social interaction, and repetitive behaviors (62). The pathophysiology of ASD is largely influenced by strong genetic components throughout the early stages of development (62). The Simmons Foundation Autism Research Initiative (SFARI) gene database (https://gene.sfari.org) reports more than 1,000 candidate genes and copy number variations (CNVs) loci associated with ASD. Importantly, many of these ASD candidate genes are implicated in synaptic formation and function (63). In fact, several molecular, cellular and functional studies of ASD experimental models suggest that synaptic abnormalities underlie ASD pathogenesis (known as synaptopathy) (64). However, the mechanisms by which these gene products induce synaptopathy and ASD symptoms are not completely understood. The *δ-catenin* gene is strongly associated with ASD according to multiple human genetic studies (20–24). Notably, the ASD-associated G34S mutation in the *δ-catenin* gene induces a loss of δ-catenin functions in excitatory synapses (24). However, how the ASD-associated δ-catenin missense mutation causes the loss of f δ-catenin functions to contribute to ASD synaptopathy remains unclear. Here, we use neuroblastoma cells to discover that the ASD-associated δ-catenin G34S mutation creates an additional phosphorylation site for GSK3β to enhance δ-catenin degradation, which is completely reversed by the reduction of GSK3β activity. Additionally, using two mouse models, δ-catenin G34S knockin and δ-catenin KO, we further identify that the δ-catenin deficiency alters cortical glutamatergic E/I at the cellular levels and disrupts social behaviors. Moreover, inhibition of GSK3β activity can reverse δ-catenin G34S-induced abnormal glutamatergic E/I and social deficits. Finally, we show that δ-catenin is required for lithium-induced restoration of normal social behavior in δ-catenin G34S mice. As the expression of δ-catenin is highly correlated with other ASD-risk genes that are involved in synaptic structure and function (31, 36–38), our findings can help us to better understand ASD synaptopathy.

In the previous works addressing a loss of δ-catenin functions, two mouse lines of δ-catenin deficiency were used. The first known mouse model contains a truncated form of δ-catenin protein (δ-catenin N-term mice) (35), making it unable to address comprehensive loss-of-function effects. Furthermore, these mice have not been used to investigate the role of δ-catenin in social behaviors. The second mouse model that has disruption in the *δ-catenin* gene has showed impaired social behaviors (65). The study, however, has not confirmed that δ-catenin protein expression is completely absent. Furthermore, this study exclusively used male mice, obviating the possibility of addressing sex disparities, which are common in many mental illnesses that are accompanied by social dysfunction (66). In contrast, we examine female and male mice separately and finds that our new δ-catenin KO female and male mice contain no full-length or truncated δ-catenin protein expression (**Fig. 5a, 5b, and S5c**) and exhibit social dysfunction (**Fig. 6a and 6b**). Therefore, employing δ-catenin G34S knockin and δ-catenin KO mouse lines allows us to examine the δ-catenin-mediated neural mechanisms underlying social behaviors in greater detail.

There may be many different factors that make a child more likely to have ASD, including environmental and genetic factors (67, 68). The interaction of both environmental and genetic risk factors for ASD is an important area of investigation as the complementary contribution of the two factors may enhance the etiologic strength in ASD pathogenesis (69). However, the molecular link of environmental and genetic risk factors for ASD has not been fully understood (70). In addition to genetic mutations, exposure to exogenous drugs during pregnancy may also cause a loss of δ-catenin functions in the fetus and interfere with brain development and maturation. One such exogenous chemical is valproic acid (VPA). Children born to mothers who were treated with VPA during pregnancy exhibit an increased incidence of neurological diseases, including ASD (71–76). VPA treatment disrupts synapse formation in developing neurons (77), as is also evident in neurons lacking δ-catenin expression (34, 78). In fact, it has been shown that VPA treatment strongly reduces δ-catenin mRNA levels in human embryonic/induced pluripotent stem (ES/iPS) cell-derived glutamatergic neurons (77). Moreover, we find that VPA-induced disruptions in synapse development are reversed by δ-catenin expression in cultured neurons (Data not shown). VPA-induced ASD pathology can thus be mediated by the loss of δ-catenin functions. This suggests that the loss of δ-catenin functions can be the common ASD pathological target arising from both genetic and environmental factors.

Another important aspect in the current study involves a shared etiology of ASD with other related disorders. For example, the Cri-du-chat syndrome results from variable hemizygous deletions in the short arm of chromosome 5p, including the *δ-catenin* gene. The clinical features of the Cri-du-chat syndrome comprise growth delay, severe mental retardation, and speech delay (79, 80). In those individuals with the syndrome who have mild or no obvious intellectual disability, the deletion often does not include the *δ-catenin* gene (81). Moreover, individuals carrying a partial deletion and a partial duplication of the *δ-catenin* gene exhibit the Cri-du-chat syndrome, but only mild cognitive disability (82). This suggests the *δ-catenin* gene plays crucial roles in etiology of the syndrome. Moreover, approximately 40% of individuals with this syndrome show some autistic-like characteristics (83). This is considerably higher than expected in the general population (83). Importantly, genetic studies identify a loss of δ-catenin functions as a significant factor for both the Cri-du-chat syndrome and ASD (21, 84–86). Moreover, altered δ-catenin functions have been implicated in many other neurological disorders, including cerebral palsy, schizophrenia, anxiety disorders, Alzheimer’s disease, and attention deficit hyperactivity disorder (87–91). Therefore, our findings likely lead to an understanding of shared etiology and pathophysiology among these diseases.

We show that the loss of δ-catenin functions by the G34S mutation and KO has the opposite effects on glutamatergic activity in cortical excitatory and inhibitory cells (**Fig. 3c, 3d, 5c, and 5d**). Consistently, a recent study demonstrates that δ-catenin deficiency increases the E/I ratio and intrinsic excitability by decreasing inhibitory synaptic activity in cortical neurons from young mice (31). However, it remains unclear how the loss of δ-catenin functions differentially affects cellular excitation and inhibition. GluA2-containing AMPARs are Ca^2+^ impermeable, whereas GluA2-lacking AMPARs are Ca^2+^ permeable (92). Although cortical excitatory neurons have no basal Ca^2+^-permeable AMPAR expression, cortical inhibitory interneurons contain GluA2-lacking Ca^2+^ permeable AMPARs (93). Therefore, δ-catenin G34S and KO-induced reduction of GluA2 in excitatory cells may promote the expression of highly conductive GluA2-lacking Ca^2+^-permeable AMPARs, which contributes to higher glutamatergic and neuronal activity. Conversely, in δ-catenin G34S and KO inhibitory interneurons, GluA1/2 or GluA2/3 heteromeric AMPARs are likely decreased, which leads to an overall decrease in glutamatergic and neuronal activity. This suggests that while δ-catenin deficiency lowers synaptic GluA2 levels in cortical neurons, it has the opposite effects on glutamatergic and neuronal activity in excitatory and inhibitory cells, potentially altering the E/I balance. Importantly, cortical network activity is known to regulate social behavior and primarily dependent on the balance between cellular E/I (94). Consistently, a large body of studies show that the medial prefrontal cortex (mPFC) activity at the cellular and network levels is strongly involved in proper social behaviors (11, 13, 14, 19, 95–97). In fact, disruptions in connectivity or E/I within the mPFC neural network have been implicated in individuals with ASD, as well as in mouse models of ASD (11–19). Therefore, maintaining the proper balance of cellular E/I and neural network activity in the mPFC is essential to ensure normal social behaviors. Importantly, glutamatergic inputs to prefrontal excitatory and inhibitory neurons from the contralateral mPFC, ventral hippocampus, basolateral amygdala, and mediodorsal thalamus are important for social behaviors (6–10). This suggests that cortical δ-catenin regulates postsynaptic glutamatergic activity in prefrontal neurons to adjust cortical network computations to control social behaviors. However, it is unknown how postsynaptic glutamatergic inputs to mPFC neurons regulate mPFC network computations to control social behaviors. Therefore, the current work is significant because we discover how δ-catenin regulates postsynaptic glutamatergic activity in cortical neurons and social behaviors.

Pharmacological treatment of ASD is challenging, due in large part to the heterogeneity in the presentation of ASD (98). Existing therapies aim to treat specific symptoms, rather than address the basic underlying etiologies (99). Thus, new areas of research focusing on therapeutic agents that directly target the underlying pathology of ASD are an important current and future need. Recent studies suggest that GSK3β activity is strongly implicated in the pathophysiology of ASD (39–46). However, extensive studies have yielded inconsistent data on the role of GSK3β activity in the molecular and behavioral effects. For example, elevated GSK3β activity is responsible for the ASD-related phenotypes in the mouse model of fragile X mental retardation (FMR) (39–42). Conversely, inactivation of GSK3β is associated with the ASD-related phenotypes found in deletion of the *phosphatase and tensin homolog on chromosome ten (PTEN)* gene in mice (43). Nonetheless, our study identifies GSK3β as the target to complete the synaptic δ-catenin regulatory pathway and recognizes GSK3β as a new therapeutic target for limiting the loss of δ-catenin function-induced synaptopathy in ASD. Moreover, we use lithium to demonstrate inhibition of GSK3β activity as a therapeutic target. Lithium reduces GSK3β activity by elevating the Akt-dependent phosphorylation of the GSK3β autoinhibitory residue, serine 9 (100) or by competing with magnesium for enzyme binding (101). Interestingly, there is strong evidence that lithium acts as a mood stabilizer through inhibition of GSK3β (102, 103).

Furthermore, one therapeutic action of lithium is the ability to regulate protein stability, in particular synaptic proteins (47, 104). Significantly, we have previously reported that lithium reduces GSK3β activity and stabilizes synaptic δ-catenin, thus elevating the δ-catenin complex consisting of AMPARs and synaptic activity in hippocampal synapses (29). Additionally, the current study demonstrate that lithium can elevate synaptic δ-catenin and GluA2 levels in the cortex and enhances social behaviors in mice, which requires δ-catenin expression. Indeed, lithium administration to 30 children and adolescents diagnosed with ASD improves the symptomatology in 43% of patients (105). Therefore, an increase in δ-catenin levels at synapses by the reduction of GSK3β activity stabilizes synaptic AMPARs and spine integrity, which could be beneficial for ASD. Therefore, our work further allows us to test a clinically applicable therapeutic protocol for ASD.

## Methods and Materials

### Animals

All mice were bred in the animal facility at Colorado State University (CSU). Animals were housed under 12:12 hour light/dark cycle. 3-month-old male and female mice were used in the current study. CSU’s Institutional Animal Care and Use Committee (IACUC) reviewed and approved the animal care and protocol (3408).

δ-catenin G34S mice (RRID:MMRRC_050621-UCD) were generated by Dr. Jyothi Arikkath at UNMC Mouse Genome Engineering Core using the CRISPR-Cas9 technique (54). Human and mouse δ-catenin proteins both contain glycine 34, according to amino acid sequence analysis (**Fig. S4a**). The CRISPR-Cas9 reagents (sgRNA, Cas9 mRNA and the donor DNA template) were injected into zygotes derived from C57BL6/J mice. The sgRNA sequence was 5’-CCATTGGAGGTGTTTAA/GCC*TGG*-3’ (the cleavage site is shown by “/” and the PAM sequence is in italics). The sgRNA is in antisense orientation, and it cleaved immediately downstream of the codon GGC (glycine 34) of the *δ-catenin* gene, yielding a double-strand break. Non-homologous end joining was used to repair the breaks, which resulted in the G34S mutation being inserted into the *δ-catenin* gene *in vivo.* The offspring were genotyped using PCR-RFLP followed by sequencing. The founder mouse was crossed with WT C57Bl6/J mice to create heterozygous δ-catenin G34S mice, which were subsequently used to make homozygous G34S animals. For genotyping, mouse tail DNA fragments were amplified by PCR using the following primers: F: 5’-GTCCAGACAGTTTCTAACTTTCATTC-3’ and R: 5’-ATGTTTCAGTGTTCCAGAGGAACAGC-3’. PCR genotyping followed by HindIII digestion yielded 250 bp and 150 bp bands for homozygous δ-catenin G34S mice (G34S), 400 bp, 250 bp, and 150 bp bands for heterozygous δ-catenin G34S mice (HET), and a 400 bp band for WT (**Fig. S4b**). Each genotype was thus identified using PCR genotyping followed by HindIII digestion (**Fig. S4c**).

The δ-catenin KO mouse was developed in collaboration with Cyagen Biosciences Inc. utilizing the CRISPR-Cas9 technique (54). Cas9 mRNA and two single guide RNAs (sgRNAs) were microinjected into the C57Bl6/J zygotes, where sgRNAs directed Cas9 endonuclease to cleave within intron 1 and intron 2 of the mouse *δ-catenin* gene (**Fig. S5a**), yielding a double-strand break, which removed the exon 2 that contained the ATG initiation codon. The sgRNA sequences were 5’-GAGGGGGGAGAGGTCCGTTC<u>TGG</u>-3’ and 5’-CTTCTTTAGGTGAACGTTGA<u>TGG</u>-3’ (the protospacer adjacent motif (PAM) sequences are underlined). Such breaks were repaired by non-homologous end joining, resulting in disruption of the *δ-catenin* gene expression *in vivo.* Cyagen Biosciences Inc. identified the optimal sequences for guide RNAs using the C57Bl6/J database to minimize off-target effects. sgRNAs that contain one or two-nucleotide mismatches to the DNA target in the 20-mer targeting region of the sgRNA do not function in the CRISPR-Cas9 system (106). We were unable to locate one or two base-mismatched sequences in the database for our sgRNAs, indicating that that δ-catenin KO was most likely successfully introduced without off-target effects. The founder mouse was crossed with WT C57Bl6/J mice to create heterozygous δ-catenin KO mice, which were subsequently used to make homozygous KO animals in the C57Bl6/J background. Each genotype was identified using PCR genotyping (**Fig. S5b**). For genotyping, mouse tail DNA fragments were amplified by PCR using the following primers: F1: 5’-TTAATCCATCGTGCTGCCAGTG-3’, R1: 5’-CTCATCATAAGAAACACCTGGAAGG-3’, and R2: 5’-CTCATACACATTCAGATTTAACCC-3’. PCR genotyping yielded a 675 bp band for homozygous δ-catenin KO mice (KO), 440 bp and 675 bp bands for heterozygous δ-catenin KO mice (HET), and a 440 bp band for WT (**Fig. S5b**). The animals used in the proposed research have been fully crossed to C57Bl6/J mice for more than eight generations and will be maintained on C57Bl6/J background. By immunoblotting whole brain tissues, we confirmed that our new δ-catenin KO mice did not express any full-length or truncated δ-catenin protein (**Fig. S5c**).

Lithium treatment in animals was conducted as described previously (29). Lithium animals received standard chow containing 0.17% w/w lithium carbonate (Harlan Teklad, Madison, WI) *ad libitum* for 7 days. Control animals were fed with standard chow.

### DNA Cloning

pSinRep5-WT δ-catenin and pSinRep5-G34S δ-catenin plasmids were gifts from Dr. Edward Ziff (New York University Langone Health). HA-tagged WT δ-catenin and G34S δ-catenin were cloned into the mammalian expression vector, pcDNA3.1. The QuikChange XL Site-Directed Mutagenesis Kit (Agilent Technologies) was used to generate G34A and G34D mutations from the WT δ-catenin plasmid. The following primers were used with the bolded regions being where the mutations were made to exchange the glycine (GGC): G34A primers 5’-GCTCCTTGAGCCCA***GCC***TTAAACACCTCCAA-3’ and 5’-TTGGAGGTGTTTAAGGCTGGGCTCAAGGAGC-3’, and the G34D primers 5’-CAGCTCCTTGAGCCCA***GAC***TTAAACACCTCCAATG-3’ and 5’-CATTGGAGGTGTTTAAGTCTGGGCTCAAGGAGCTG-3’.

### Reagents

Proteasome inhibitor MG132 (Allfa Aesar) was used at 10 μM to treat SH-SY5Y cells 4 hours before collecting whole cell lysates. 2 mM lithium chloride (LiCl) (Sigma-Aldrich) was used to inhibit GSK3β activity in SH-SY5Y cells and cultured cortical neurons, and 2 mM sodium chloride (NaCl) (Sigma-Aldrich) was used as a control. 2 μM tetrodotoxin (TTX) (Abcam) was used to block spontaneous Ca^2+^ activity in cultured cortical neurons. 1 μM 4-methoxy-7-nitroindolinyl (MNI)-caged L-glutamate (Tocris Bioscience) was added to the culture media for glutamate uncaging.

### Human neuroblastoma cell (SH-SY5Y) culture and transfection

SH-SY5Y cells were grown in DMEM medium with fetal bovine serum (Life Technologies) and 1% penicillin/streptomycin (Life Technologies). 500,000 cells were plated in 6-well dishes and 1 μg of DNA was transfected when cells reached 75-85% confluency with Lipofectamine 2000 (Life Technologies) according to the manufacturer’s instructions. 25 nM control siRNA ON-TARGETplus non-targeting pool (Dharmacon siRNA solution) and 25 nM SMARTpool ON-TARGETplus human GSK3β siRNA (Dharmacon siRNA solution) were transfected with 1 μg of DNA when cells reached 50% confluency and cell lysates were collected 72 hours later. Cells used for each experiment were from more than three independently prepared cultures.

### Primary hippocampal neuronal culture

Postnatal day 0 (P0) male and female pups from WT, δ-catenin G34S or KO mice were used to produce mouse cortical neuron cultures as shown previously (107–109). Heterozygous G34S or KO mice were used to breed for generating each genotype. PCR was performed to identify each genotype before preparing cultures. Cortical tissues were isolated from P0 mouse brains and digested with 10 U/mL papain (Worthington Biochemical Corp. Lakewood, NJ). Mouse cortical neurons were plated on following poly lysine-coated glass bottom dishes (500,000 cells) for Ca^2+^ imaging with glutamate uncaging. Cells were grown in Neurobasal Medium without phenol red (Thermo Fisher Scientific) with B27 supplement (Thermo Fisher Scientific), 0.5 mM Glutamax (Thermo Fisher Scientific), and 1% penicillin/streptomycin (Thermo Fisher Scientific).

### Immunoblotting

Immunoblotting was performed as described previously (29, 107–120). The protein concentration in the samples was determined by a BCA protein assay kit (Thermo Fisher Scientific). Equal amounts of protein samples were loaded on 10% glycine-SDS-PAGE gel. The gels were transferred to nitrocellulose membranes. The membranes were blocked (5% powdered milk) for 1 hour at room temperature, followed by overnight incubation with the primary antibodies at 4°C. The primary antibodies consisted of anti-δ-catenin (BD Biosciences, 1:1000 and Abcam, 1:1000), anti-GluA1 (Millipore, 1:2000), anti-GluA2 (Abcam, 1:2000), anti-pGSK3β (Cell Signaling Technology, 1:1000), and anti-actin (Abcam, 1:2000) antibodies. Membranes were subsequently incubated by secondary antibodies for 1 hour at room temperature and developed with Enhanced Chemiluminescence (ECL) (Thermo Fisher Scientific, Waltham, MA). Protein bands were quantified using ImageJ (https://imagei.nih.gov/ij/).

### GCaMP Ca^2+^ Imaging with glutamate uncaging

We carried out Ca^2+^ imaging with glutamate uncaging in cultured mouse cortical neurons to determine glutamatergic activity as described previously (121). For Ca^2+^ imaging, a genetically encoded Ca^2+^ indicator, GCaMP, was used. When AAVs of the same serotype are co-infected, many neurons are transduced by both viruses (122). We thus co-infected AAVs expressing CamK2a-Cre (Addgene# 105558-AAV1), pENN.AAV.CamKII 0.4.Cre.SV40 was a gift from James M. Wilson (Addgene plasmid # 105558; http://n2t.net/addgene:105558; RRID:Addgene_105558), and Cre-dependent GCaMP7s (Addgene# 104495-AAV1) (123), pGP-AAV-CAG-FLEX-jGCaMP7s-WPRE was a gift from Douglas Kim & GENIE Project (Addgene plasmid # 104495; http://n2t.net/addgene:104495; RRID:Addgene_104495), in 4 days *in vitro* (DIV) neurons and imaged 12-13 DIV excitatory neurons. In addition, AAV expressing GCaMP6f under the control of the GABAergic neuron-specific enhancer of the mouse *Dlx* (mDlx) gene (Addgene# 83899-AAV1) (124), pAAV-mDlx-GCaMP6f-Fishell-2 was a gift from Gordon Fishell (Addgene plasmid # 83899-AAV1; http://n2t.net/addgene:83899; RRID:Addgene_83899), was infected in 4 DIV neurons and imaged 12-13 DIV inhibitory interneurons. Glass-bottom dishes were mounted on a temperature-controlled stage on an Olympus IX73 microscope and maintained at 37°C and 5% CO_2_ using a Tokai-Hit heating stage and digital temperature and humidity controller. For glutamate uncaging, 1 μM 4-methoxy-7-nitroindolinyl (MNI)-caged L-glutamate was added to the culture media, and epi-illumination photolysis (390 nm, 0.12 mW/mm^2^, 1 ms) was used. 2 μM TTX was added to prevent action potential-dependent network activity. A baseline average of 20 frames (10 ms exposure for GCaMP7s and 50 ms exposure for GCaMP6f exposure) (F_0_) were captured prior to glutamate uncaging, and 50 more frames were obtained after single photostimulation. The fractional change in fluorescence intensity relative to baseline (ΔF/F0) was calculated. The average peak amplitude in the control group was used to normalize the peak amplitude in each cell. The control group’s average peak amplitude was compared to the experimental groups’ average.

### Three-chamber test

To determine social behaviors, a three-chamber test was performed with a modification of the previously described method (118). Before the test session, the subject mouse was placed in the center chamber and allowed to habituate for 5 minutes. In the sociability test (10 minutes), an unfamiliar mouse (stranger 1) was placed in one side chamber under an inverted stainless-steel wire cup that allowed olfactory, visual, auditory, and tactile contacts, and an empty cup (a novel object) was placed in the opposite side chamber. During the social novelty test (10 minutes), a new unfamiliar mouse (stranger 2) was placed under a wire cup in the opposite side chamber that had been empty during the sociability phase. The subject mouse was allowed to explore freely in all 3 chambers during the tests. The behavior was recorded using a camera mounted overhead. The test animals’ interaction with strangers was determined by the reciprocal sniffing time as described previously (125, 126). The test animals’ interaction with a novel object was determined by the sniffing time that was defined as each instance in which a test mouse’s nose was oriented in a 10-degree head orientation and comes within 2 cm toward a mouse or a wire cup as described previously (14). The discrimination index was calculated as (Total reciprocal sniffing time for stranger 1 – Total sniffing time for the object) / (Total reciprocal sniffing time for stranger 1 + Total sniffing time for the object) for the sociability test and (Total reciprocal sniffing time for stranger 2 – Total reciprocal sniffing time for stranger 1) / (Total reciprocal sniffing time for stranger 2 + Total reciprocal sniffing time for stranger 1) for the social novelty test. We confirmed the reciprocal sniffing time manually and blindly by two different investigators.

### Buried food assay

The buried food assay was performed as described previously (111). Several days before the test, 2-3 Froot Loops (Kellogg’s) were placed in each cage overnight to confirm that the food is palatable to the mice. Mice that did not consume the Froot Loops were omitted from the test. Mice were food-deprived for 24 hours prior to the test. 2 Froot Loops were placed in a clean cage and buried under fresh bedding prior to placing the mice in the cage. Mice were allowed 20 minutes to explore the cage searching for the hidden food and the latency to find and begin to nibble on the food was recorded. Mice that did not find the food after 20 minutes were excluded from the results. The latency was blindly scored by two different investigators.

### Open field test

We measured locomotor activity and anxiety-like behavior using the open field test as carried out previously (118). The test mouse was first placed in the center of the open field chamber (40 W x 40 L x 40 H cm) for 5 minutes. Animals were then allowed to explore the chamber for 20 minutes. A 20 x 20 cm center square was defined as the inside zone. The behavior was recorded by a video camera. Data were analyzed using the ANY-maze tracking program to acquire total traveled distance (locomotor activity) and time spent outside and inside (anxiety-like behavior).

### Statistical analysis

The Franklin A. Graybill Statistical Laboratory at CSU has been consulted for statistical analysis in the current study, including sample size determination, randomization, experiment conception and design, data analysis, and interpretation. We used the GraphPad Prism 9 software to determine statistical significance (set at *p* < 0.05). Grouped results of single comparisons were analyzed using the unpaired two-tailed Student’s t-test. Differences between multiple groups were assessed by Two-way analysis of variance (ANOVA) with the Tukey test or nonparametric Kruskal-Wallis test with the Dunn’s test. The graphs were presented as mean ± Standard Deviation (SD).

## Acknowledgments

We thank members of the Kim laboratory for their generous support. We appreciate thoughtful suggestion from Dr. Edward Ziff. This work is supported by Student Experiential Learning Grants from CSU, the Jerome Lejeune Foundation, the NIH/NCATS Colorado CTSA Grant (UL1 TR002535), the Boettcher Foundation’s Webb-Waring Biomedical Research Program, and the NIH grant (1R03AG072102). This work was prepared while Jyothi Arikkath was employed at the University of Nebraska Medical Center. The opinions expressed in this article are the author’s own and do not reflect the views of the National Institutes of Health, the Department of Health and Human Services, or the United States Government.

---

**Supplementary Figure 1.**
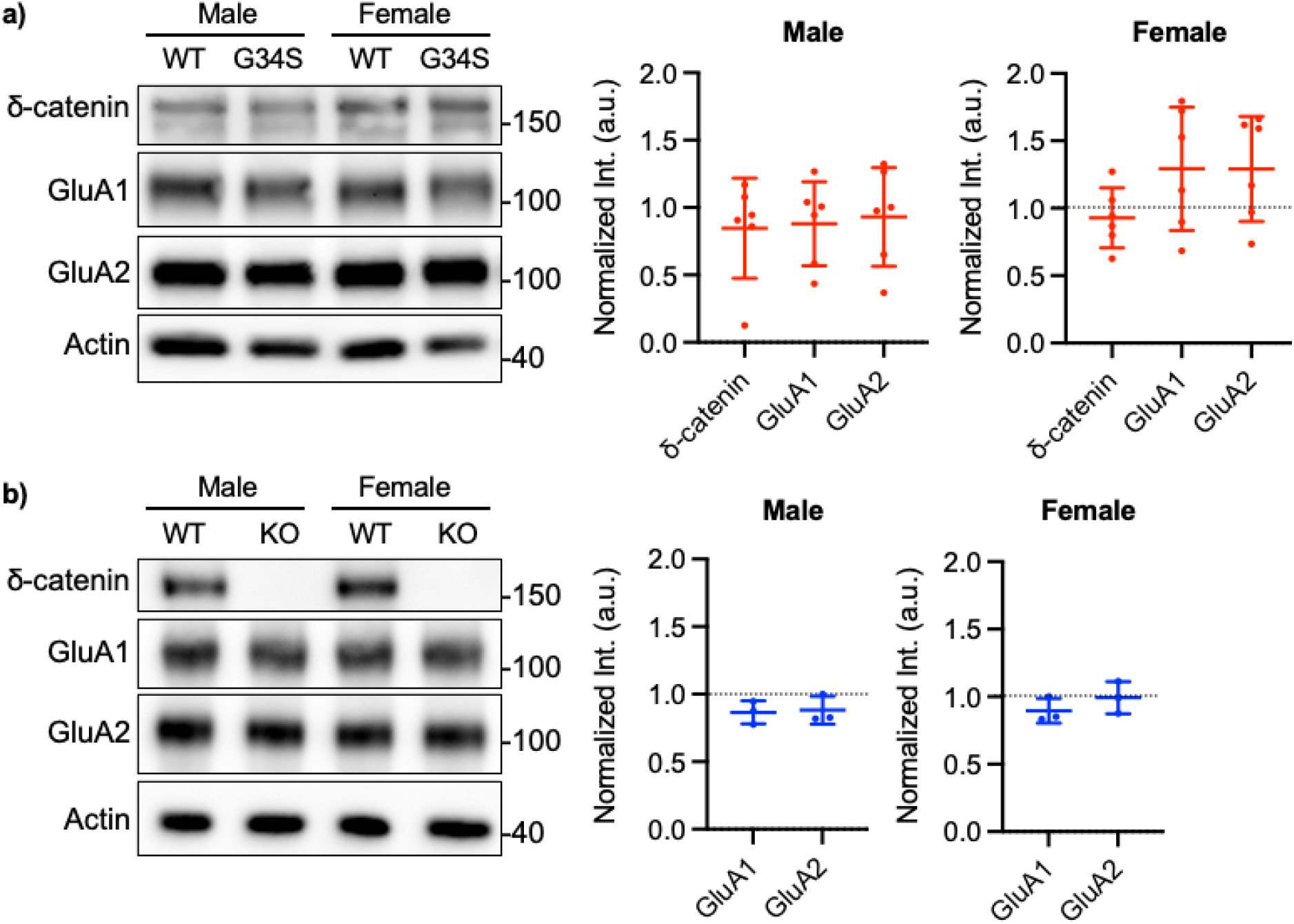
Synaptic δ-catenin and AMPAR levels in the hippocampus of δ-catenin G34S and KO mice. **a)** Representative immunoblots and summary graphs of normalized δ-catenin, GluA1, and GluA2 levels in the male and female hippocampal PSD fractions from WT and δ-catenin G34S showing no difference in synaptic δ-catenin and AMPAR levels between the WT and G34S hippocampus (n = 6 immunoblots from 3 mice in each condition, Kruskal-Wallis test with the Dunn’s test). **b)** Representative immunoblots and summary graphs of normalized GluA1 and GluA2 levels in the male and female hippocampal PSD fractions from WT and δ-catenin KO showing no difference in synaptic δ-catenin and AMPAR levels between the WT and KO hippocampus (n = 3 immunoblots from 3 mice in each condition, Kruskal-Wallis test with the Dunn’s test). No δ-catenin expression was found in the KO hippocampus. The position of molecular mass markers (kDa) is shown on the right of the blots.

**Supplementary Figure 2.**
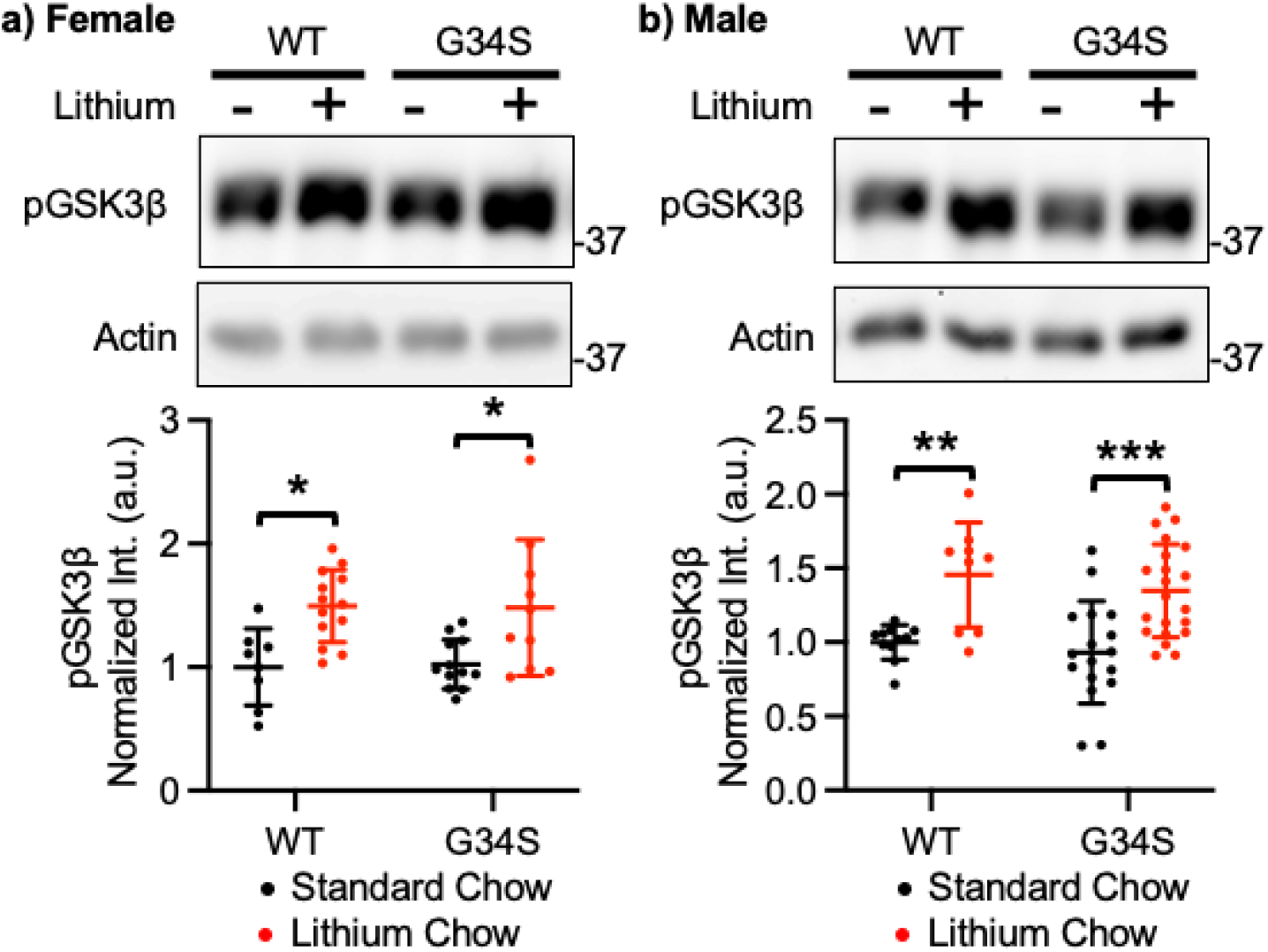
Lithium treatment significantly reduces *in vivo* GSK3β activity. Representative immunoblots and summary graphs of normalized pGSK3β levels in the whole cell lysates from the cortex of standard chow (-) or lithium chow (+)-fed WT and δ-catenin G34S **a)** female (n=number of immunoblots [number of mice]. WT female + standard chow = 8 [6], WT female + lithium chow = 13 [6], G34S female + standard chow = 12 [6], and G34S female + lithium chow = 10 [6]) and **b)** male cortex (WT male + standard chow=11 [6], WT male + lithium chow = 9 [6], G34S male + standard chow = 17 [6], and G34S male + lithium chow = 21 [6]). \**p* < 0.05, \*\**p* < 0.01, and \*\*\**p* < 0.001, Two-way ANOVA with the Tukey test). The position of molecular mass markers (kDa) is shown on the right of the blots.

**Supplementary Figure 3.**
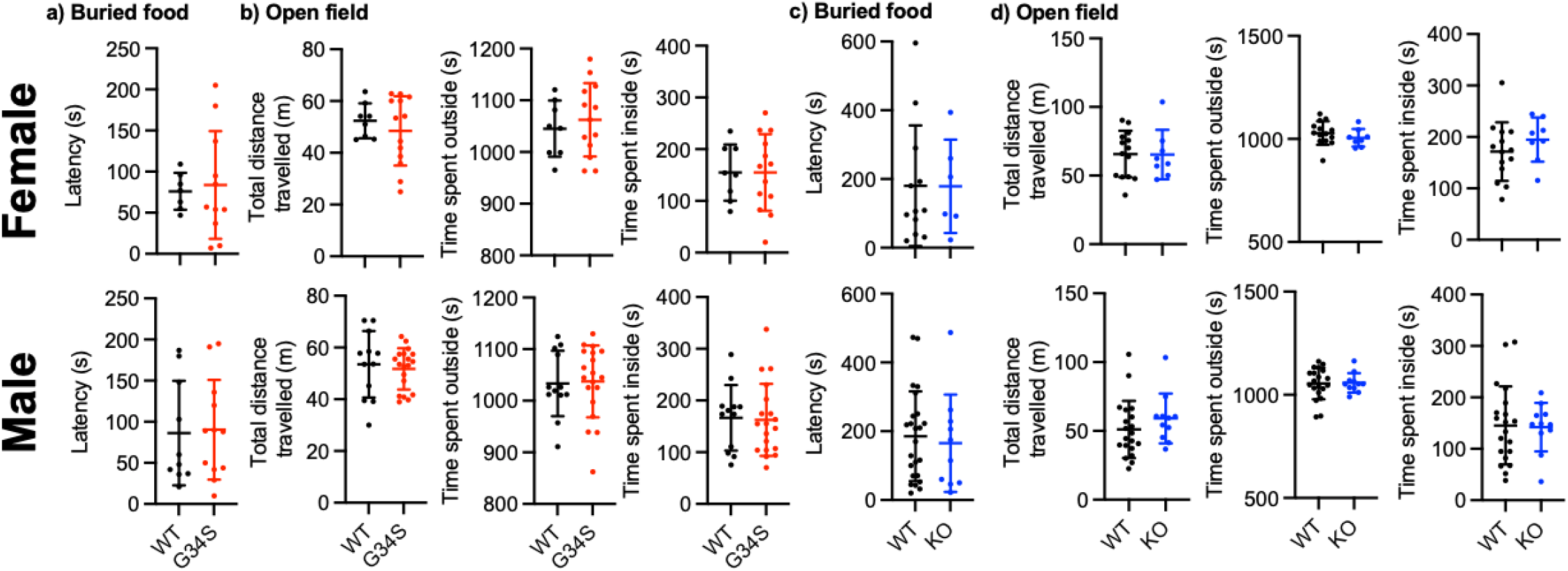
Normal olfaction, locomotor activity, and anxiety levels in δ-catenin G34S and KO mice. The results of the buried food test showing normal olfaction in δ-catenin G34S **a)** females (n = 7 WT and 11 G34S mice) and males (n = 10 WT and 12 G34S mice). **b)** The results of the open field test measuring total distance travelled and time spent outside and inside showing normal locomotor activity and no anxiety-like behavior in δ-catenin G34S females (n = 8 WT and 13 G34S mice) and males (n = 12 WT and 19 G34S mice). The results of the buried food test showing normal olfaction in δ-catenin KO **c)** females (n = 12 WT and 6 KO mice) and males (n = 24 WT and 10 KO mice). **d)** The results of the open field test measuring total distance travelled and time spent outside and inside showing normal locomotor activity and no anxiety-like behavior in δ-catenin KO females (n = 18 WT and 8 KO mice) and males (n = 20 WT and 11 KO mice). The unpaired twotailed Student’s t-test.

**Supplementary Figure 4.**
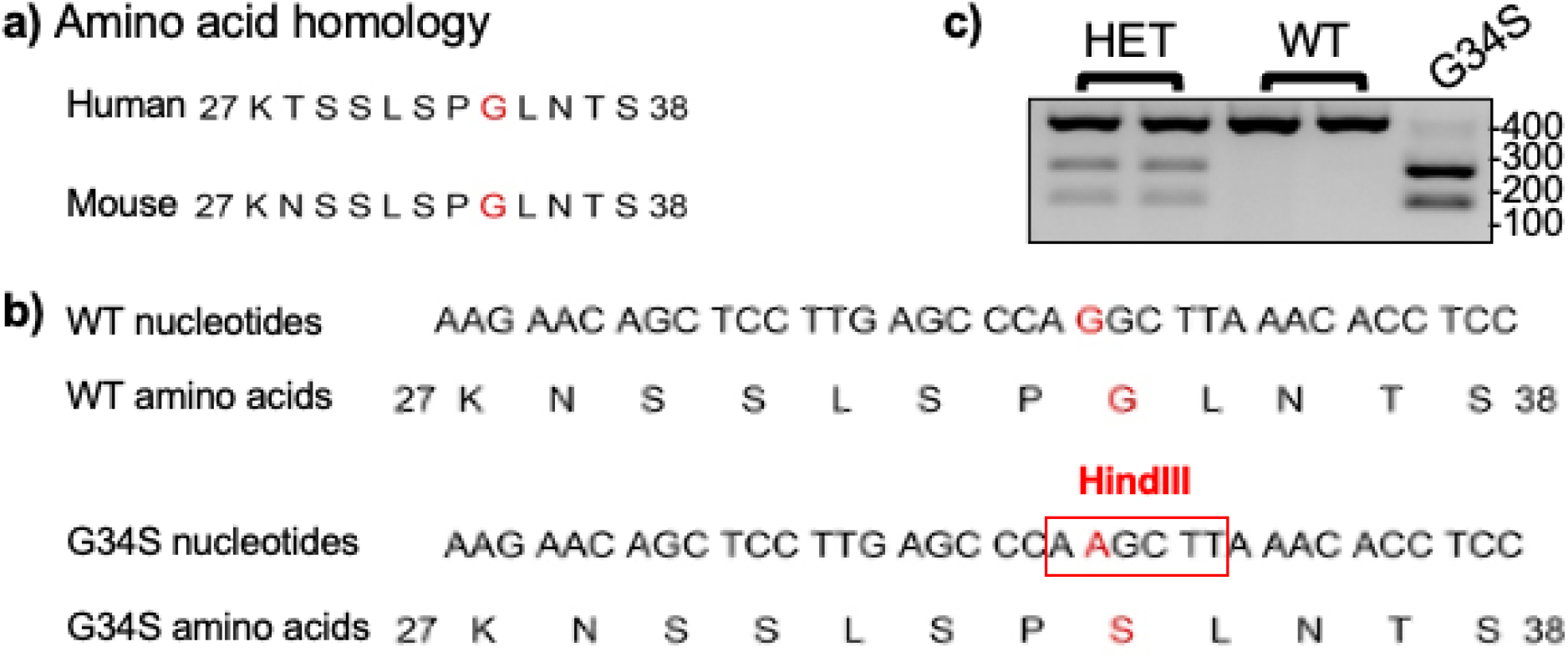
Generation of δ-catenin G34S mice. **a)** Amino acid sequence homology between human and mouse δ-catenin at indicated locus. **b)** Nucleotide sequence and corresponding amino acid sequence for WT and mutant alleles. The HindIII site is highlighted. **c)** For genotyping, mouse tail DNA fragments were amplified by PCR. PCR genotyping followed by HindIII digestion yields 250 bp and 150 bp bands for homozygous δ-catenin G34S mice (G34S), 400 bp, 250 bp, and 150 bp bands for heterozygous δ-catenin G34S mice (HET), and a 400 bp band for WT. The position of DNA size markers (bp) is shown on the right of the image.

**Supplementary Figure 5.**
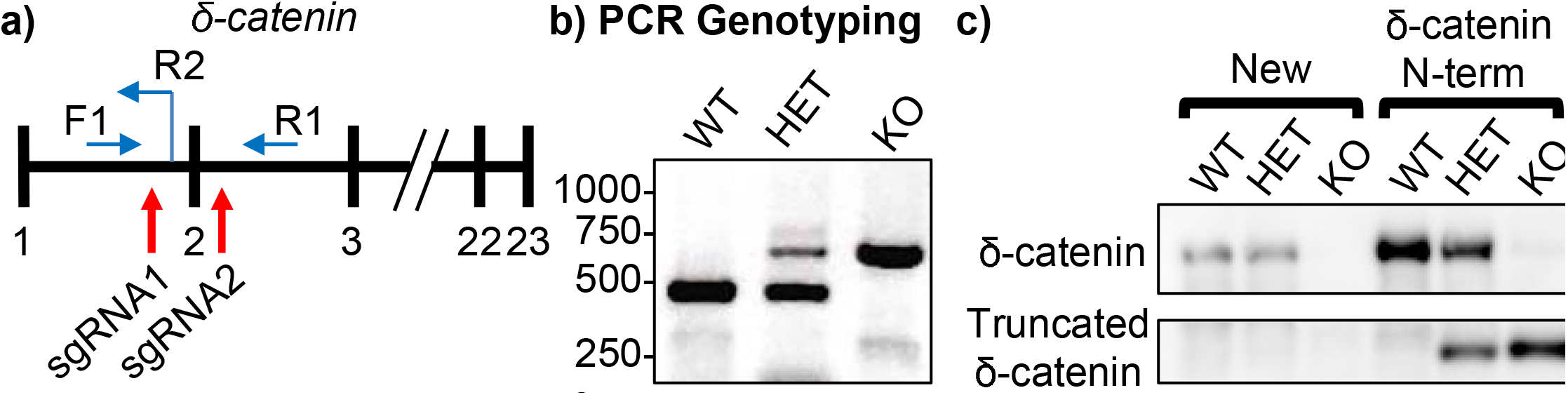
Generation of δ-catenin KO mice. **a)** The strategy of generation of δ-catenin KO mice. The numbers indicate exons, red arrows indicate sgRNA target sites, and blue arrows indicate primers used for PCR genotyping. **b)** PCR genotyping yields a 675 bp band for homozygous δ-catenin KO mice (KO), 440 bp and 675 bp bands for heterozygous δ-catenin KO mice (HET), and a 440 bp band for WT. The position of DNA size markers (bp) is shown on the left of the image. **c)** Representative immunoblot images of δ-catenin levels in total cell lysates from the cortex of WT, HET, and KO mice of newly generated δ-catenin KO (New) and δ-catenin N-term animals showing that new δ-catenin KO mice have no full-length and truncated δ-catenin. The position of molecular mass markers (kDa) is shown on the right of the blots.

## Notes

### Competing Interest Statement

The authors have declared no competing interest.

## References

1. J. Ko, Neuroanatomical Substrates of Rodent Social Behavior: The Medial Prefrontal Cortex and Its Projection Patterns. Frontiers in neural circuits 11, 41 (2017).

2. I. Rapin, R. Katzman, Neurobiology of autism. Ann Neurol 43, 7–14 (1998).

3. R. C. Abrams, L. A. Spielman, G. S. Alexopoulos, E. Klausner, Personality disorder symptoms and functioning in elderly depressed patients. Am J Geriatr Psychiatry 6, 24–30 (1998).

4. J. O. Goldberg, L. A. Schmidt, Shyness, sociability, and social dysfunction in schizophrenia. Schizophr Res 48, 343–349 (2001).

5. A. P. Association, Diagnostic and statistical manual of mental disorders (DSM-5®) (American Psychiatric Pub, 2013).

6. Q. Sun et al., Ventral Hippocampal-Prefrontal Interaction Affects Social Behavior via Parvalbumin Positive Neurons in the Medial Prefrontal Cortex. iScience 23, 100894 (2020).

7. A. C. Felix-Ortiz, A. Burgos-Robles, N. D. Bhagat, C. A. Leppla, K. M. Tye, Bidirectional modulation of anxiety-related and social behaviors by amygdala projections to the medial prefrontal cortex. Neuroscience 321, 197–209 (2016).

8. J. P. Little, A. G. Carter, Subcellular synaptic connectivity of layer 2 pyramidal neurons in the medial prefrontal cortex. J Neurosci 32, 12808–12819 (2012).

9. S. Xu et al., Neural Circuits for Social Interactions: From Microcircuits to Input-Output Circuits. Frontiers in neural circuits 15, 768294 (2021).

10. B. R. Ferguson, W. J. Gao, Thalamic Control of Cognition and Social Behavior Via Regulation of Gamma-Aminobutyric Acidergic Signaling and Excitation/Inhibition Balance in the Medial Prefrontal Cortex. Biological psychiatry 83, 657–669 (2018).

11. W. C. Huang, Y. Chen, D. T. Page, Hyperconnectivity of prefrontal cortex to amygdala projections in a mouse model of macrocephaly/autism syndrome. Nat Commun 7, 13421 (2016).

12. A. Selimbeyoglu et al., Modulation of prefrontal cortex excitation/inhibition balance rescues social behavior in CNTNAP2-deficient mice. Sci Transl Med 9(2017).

13. O. Yizhar et al., Neocortical excitation/inhibition balance in information processing and social dysfunction. Nature 477, 171–178 (2011).

14. W. Cao et al., Gamma Oscillation Dysfunction in mPFC Leads to Social Deficits in Neuroligin 3 R451C Knockin Mice. Neuron 97, 1253–1260 e1257 (2018).

15. F. Abell et al., The neuroanatomy of autism: a voxel-based whole brain analysis of structural scans. Neuroreport 10, 1647–1651 (1999).

16. F. Castelli, C. Frith, F. Happe, U. Frith, Autism, Asperger syndrome and brain mechanisms for the attribution of mental states to animated shapes. Brain: a journal of neurology 125, 1839–1849 (2002).

17. N. Schmitz et al., Neural correlates of executive function in autistic spectrum disorders. Biological psychiatry 59, 7–16 (2006).

18. A. C. Brumback et al., Identifying specific prefrontal neurons that contribute to autism-associated abnormalities in physiology and social behavior. Molecular psychiatry 23, 2078–2089 (2018).

19. L. Liu et al., Cell type-differential modulation of prefrontal cortical GABAergic interneurons on low gamma rhythm and social interaction. Sci Adv 6, eaay4073 (2020).

20. I. O. Tuncay et al., Analysis of recent shared ancestry in a familial cohort identifies coding and noncoding autism spectrum disorder variants. NPJ Genom Med 7, 13 (2022).

21. D. E. Miller, A. Squire, J. T. Bennett, A child with autism, behavioral issues, and dysmorphic features found to have a tandem duplication within CTNND2 by mate-pair sequencing. Am J Med Genet A 10.1002/ajmg.a.61442 (2019).

22. H. Guo et al., Inherited and multiple de novo mutations in autism/developmental delay risk genes suggest a multifactorial model. Mol Autism 9, 64 (2018).

23. T. Wang et al., De novo genic mutations among a Chinese autism spectrum disorder cohort. Nat Commun 7, 13316 (2016).

24. T. N. Turner et al., Loss of delta-catenin function in severe autism. Nature 520, 51–56 (2015).

25. J. B. Silverman et al., Synaptic anchorage of AMPA receptors by cadherins through neural plakophilin-related arm protein AMPA receptor-binding protein complexes. J Neurosci 27, 8505–8516 (2007).

26. M. Peifer, S. Berg, A. B. Reynolds, A repeating amino acid motif shared by proteins with diverse cellular roles. Cell 76, 789–791 (1994).

27. M. Takeichi, Cadherins: key molecules for selective cell-cell adhesion. IARC Sci Publ, 76–79 (1988).

28. J. Gilbert, H. Y. Man, The X-Linked Autism Protein KIAA2022/KIDLIA Regulates Neurite Outgrowth via N-Cadherin and delta-Catenin Signaling. eNeuro 3 (2016).

29. M. Farooq et al., Lithium increases synaptic GluA2 in hippocampal neurons by elevating the delta-catenin protein. Neuropharmacology 10.1016/j.neuropharm.2016.10.025 (2016).

30. S. Restituito et al., Synaptic autoregulation by metalloproteases and gamma-secretase. J Neurosci 31, 12083–12093 (2011).

31. N. Assendorp et al., CTNND2 moderates neuronal excitation and links human evolution to prolonged synaptic development in the neocortex. bioRxiv 10.1101/2022.09.13.507776, 2022.2009.2013.507776 (2022).

32. K. S. Kosik, C. P. Donahue, I. Israely, X. Liu, T. Ochiishi, Delta-catenin at the synaptic-adherens junction. Trends Cell Biol 15, 172–178 (2005).

33. L. Yuan, E. Seong, J. L. Beuscher, J. Arikkath, delta-Catenin Regulates Spine Architecture via Cadherin and PDZ-dependent Interactions. The Journal of biological chemistry 290, 10947–10957 (2015).

34. C. Matter, M. Pribadi, X. Liu, J. T. Trachtenberg, Delta-catenin is required for the maintenance of neural structure and function in mature cortex in vivo. Neuron 64, 320–327 (2009).

35. I. Israely et al., Deletion of the neuron-specific protein delta-catenin leads to severe cognitive and synaptic dysfunction. Current biology: CB 14, 1657–1663 (2004).

36. S. Ramanathan et al., A case of autism with an interstitial deletion on 4q leading to hemizygosity for genes encoding for glutamine and glycine neurotransmitter receptor sub-units (AMPA 2, GLRA3, GLRB) and neuropeptide receptors NPY1R, NPY5R. BMC Med Genet 5, 10 (2004).

37. A. El-Amraoui, C. Petit, Cadherins as targets for genetic diseases. Cold Spring Harb Perspect Biol 2, a003095 (2010).

38. R. Mejias et al., Gain-of-function glutamate receptor interacting protein 1 variants alter GluA2 recycling and surface distribution in patients with autism. Proc Natl Acad Sci U S A 108, 4920–4925 (2011).

39. W. W. Min et al., Elevated glycogen synthase kinase-3 activity in Fragile X mice: key metabolic regulator with evidence for treatment potential. Neuropharmacology 56, 463–472 (2009).

40. C. J. Yuskaitis et al., Lithium ameliorates altered glycogen synthase kinase-3 and behavior in a mouse model of fragile X syndrome. Biochemical pharmacology 79, 632–646 (2010).

41. E. J. McManus et al., Role that phosphorylation of GSK3 plays in insulin and Wnt signalling defined by knockin analysis. Embo J 24, 1571–1583 (2005).

42. M. A. Mines, C. J. Yuskaitis, M. K. King, E. Beurel, R. S. Jope, GSK3 influences social preference and anxiety-related behaviors during social interaction in a mouse model of fragile X syndrome and autism. PLoS One 5, e9706 (2010).

43. C. H. Kwon et al., Pten regulates neuronal arborization and social interaction in mice. Neuron 50, 377–388 (2006).

44. Y. Mao et al., Disrupted in schizophrenia 1 regulates neuronal progenitor proliferation via modulation of GSK3beta/beta-catenin signaling. Cell 136, 1017–1031 (2009).

45. W. Y. Kim, W. D. Snider, Functions of GSK-3 Signaling in Development of the Nervous System. Frontiers in molecular neuroscience 4, 44 (2011).

46. X. Wu et al., Lithium ameliorates autistic-like behaviors induced by neonatal isolation in rats. Front Behav Neurosci 8, 234 (2014).

47. O. O’Leary, Y. Nolan, Glycogen synthase kinase-3 as a therapeutic target for cognitive dysfunction in neuropsychiatric disorders. CNS Drugs 29, 1–15 (2015).

48. S. Bareiss, K. Kim, Q. Lu, Delta-catenin/NPRAP: A new member of the glycogen synthase kinase- 3beta signaling complex that promotes beta-catenin turnover in neurons. Journal of neuroscience research 88, 2350–2363 (2010).

49. M. Oh et al., GSK-3 phosphorylates delta-catenin and negatively regulates its stability via ubiquitination/proteosome-mediated proteolysis. The Journal of biological chemistry 284, 28579–28589 (2009).

50. C. Yost et al., The axis-inducing activity, stability, and subcellular distribution of beta-catenin is regulated in Xenopus embryos by glycogen synthase kinase 3. Genes Dev 10, 1443–1454 (1996).

51. M. Peifer, L. M. Pai, M. Casey, Phosphorylation of the Drosophila adherens junction protein Armadillo: roles for wingless signal and zeste-white 3 kinase. Developmental biology 166, 543–556 (1994).

52. I. Celen, K. E. Ross, C. N. Arighi, C. H. Wu, Bioinformatics Knowledge Map for Analysis of Beta-Catenin Function in Cancer. PLoS One 10, e0141773 (2015).

53. Y. Xue et al., GPS 2.0, a tool to predict kinase-specific phosphorylation sites in hierarchy. Mol Cell Proteomics 7, 1598–1608 (2008).

54. D. W. Harms et al., Mouse Genome Editing Using the CRISPR/Cas System. Curr Protoc Hum Genet 83, 15 17 11-27 (2014).

55. N. Chen et al., Interaction proteomics reveals brain region-specific AMPA receptor complexes. J Proteome Res 13, 5695–5706 (2014).

56. M. Maheshwari, A. Samanta, S. K. Godavarthi, R. Mukherjee, N. R. Jana, Dysfunction of the ubiquitin ligase Ube3a may be associated with synaptic pathophysiology in a mouse model of Huntington disease. The Journal of biological chemistry 287, 29949–29957 (2012).

57. J. L. Rubenstein, M. M. Merzenich, Model of autism: increased ratio of excitation/inhibition in key neural systems. Genes Brain Behav 2, 255–267 (2003).

58. G. K. Beauchamp, K. Yamazaki, Chemical signalling in mice. Biochemical Society transactions 31, 147–151 (2003).

59. M. Yang, J. N. Crawley, Simple behavioral assessment of mouse olfaction. Current protocols in neuroscience /editorial board, Jacqueline N. Crawley… [et al.] Chapter 8, Unit 8 24 (2009).

60. A. K. Beery, D. Kaufer, Stress, social behavior, and resilience: insights from rodents. Neurobiol Stress 1, 116–127 (2015).

61. M. Yang, J. L. Silverman, J. N. Crawley, Automated three-chambered social approach task for mice. Current protocols in neuroscience /editorial board, Jacqueline N. Crawley… [et al.] Chapter 8, Unit 8 26 (2011).

62. S. Guang et al., Synaptopathology Involved in Autism Spectrum Disorder. Frontiers in cellular neuroscience 12, 470 (2018).

63. J. Gilbert, H. Y. Man, Fundamental Elements in Autism: From Neurogenesis and Neurite Growth to Synaptic Plasticity. Frontiers in cellular neuroscience 11, 359 (2017).

64. H. Won, W. Mah, E. Kim, Autism spectrum disorder causes, mechanisms, and treatments: focus on neuronal synapses. Frontiers in molecular neuroscience 6, 19 (2013).

65. X. Wang et al., Rictor is involved in Ctnnd2 deletion-induced impairment of spatial learning and memory but not autism-like behaviors. Front Biosci (Landmark Ed) 26, 335–346 (2021).

66. M. M. Wickens, D. A. Bangasser, L. A. Briand, Sex Differences in Psychiatric Disease: A Focus on the Glutamate System. Frontiers in molecular neuroscience 11, 197 (2018).

67. G. Huguet, E. Ey, T. Bourgeron, The genetic landscapes of autism spectrum disorders. Annu Rev Genomics Hum Genet 14, 191–213 (2013).

68. J. Christensen et al., Prenatal valproate exposure and risk of autism spectrum disorders and childhood autism. JAMA 309, 1696–1703 (2013).

69. D. F. Mabunga, E. L. Gonzales, J. W. Kim, K. C. Kim, C. Y. Shin, Exploring the Validity of Valproic Acid Animal Model of Autism. Exp Neurobiol 24, 285–300 (2015).

70. P. Chaste, M. Leboyer, Autism risk factors: genes, environment, and gene-environment interactions. Dialogues Clin Neurosci 14, 281–292 (2012).

71. F. I. Roullet, J. K. Lai, J. A. Foster, In utero exposure to valproic acid and autism--a current review of clinical and animal studies. Neurotoxicol Teratol 36, 47–56 (2013).

72. J. Jentink et al., Valproic acid monotherapy in pregnancy and major congenital malformations. N Engl J Med 362, 2185–2193 (2010).

73. G. Koren, A. A. Nava-Ocampo, M. E. Moretti, R. Sussman, I. Nulman, Major malformations with valproic acid. Can Fam Physician 52, 441–442, 444, 447 (2006).

74. S. J. Moore et al., A clinical study of 57 children with fetal anticonvulsant syndromes. J Med Genet 37, 489–497 (2000).

75. A. D. Rasalam et al., Characteristics of fetal anticonvulsant syndrome associated autistic disorder. Dev Med Child Neurol 47, 551–555 (2005).

76. G. Williams et al., Fetal valproate syndrome and autism: additional evidence of an association. Dev Med Child Neurol 43, 202–206 (2001).

77. S. Chanda et al., Direct Reprogramming of Human Neurons Identifies MARCKSL1 as a Pathogenic Mediator of Valproic Acid-Induced Teratogenicity. Cell Stem Cell 10.1016/j.stem.2019.04.021 (2019).

78. R. Baumert et al., Novel phospho-switch function of delta-catenin in dendrite development. The Journal of cell biology 219 (2020).

79. E. Niebuhr, The Cri du Chat syndrome: epidemiology, cytogenetics, and clinical features. Hum Genet 44, 227–275 (1978).

80. J. Overhauser, A. L. Beaudet, J. J. Wasmuth, A fine structure physical map of the short arm of chromosome 5. Am J Hum Genet 39, 562–572 (1986).

81. M. Medina, R. C. Marinescu, J. Overhauser, K. S. Kosik, Hemizygosity of delta-catenin (CTNND2) is associated with severe mental retardation in cri-du-chat syndrome. Genomics 63, 157–164 (2000).

82. J. M. Sardina, A. R. Walters, K. E. Singh, R. X. Owen, V. E. Kimonis, Amelioration of the typical cognitive phenotype in a patient with the 5pter deletion associated with Cri-du-chat syndrome in addition to a partial duplication of CTNND2. Am J Med Genet A 164A, 1761–1764 (2014).

83. J. F. Moss et al., Prevalence of autism spectrum phenomenology in Cornelia de Lange and Cri du Chat syndromes. Am J Ment Retard 113, 278–291 (2008).

84. L. A. Weiss et al., A genome-wide linkage and association scan reveals novel loci for autism. Nature 461, 802–808 (2009).

85. C. Harvard et al., A variant Cri du Chat phenotype and autism spectrum disorder in a subject with de novo cryptic microdeletions involving 5p15.2 and 3p24.3-25 detected using whole genomic array CGH. Clin Genet 67, 341–351 (2005).

86. B. Zhang et al., Multigenerational autosomal dominant inheritance of 5p chromosomal deletions. Am J Med Genet A 170, 583–593 (2016).

87. G. McMichael et al., Rare copy number variation in cerebral palsy. Eur J Hum Genet 22, 40–45 (2014).

88. T. Vrijenhoek et al., Recurrent CNVs disrupt three candidate genes in schizophrenia patients. Am J Hum Genet 83, 504–510 (2008).

89. M. G. Nivard et al., Further confirmation of the association between anxiety and CTNND2: replication in humans. Genes Brain Behav 13, 195–201 (2014).

90. G. Jun et al., delta-Catenin is genetically and biologically associated with cortical cataract and future Alzheimer-related structural and functional brain changes. PLoS One 7, e43728 (2012).

91. A. Adegbola et al., Disruption of CTNND2, encoding delta-catenin, causes a penetrant attention deficit disorder and myopia. HGG Adv 1 (2020).

92. G. H. Diering, R. L. Huganir, The AMPA Receptor Code of Synaptic Plasticity. Neuron 100, 314–329 (2018).

93. J. T. Isaac, M. C. Ashby, C. J. McBain, The role of the GluR2 subunit in AMPA receptor function and synaptic plasticity. Neuron 54, 859–871 (2007).

94. B. R. Ferguson, W. J. Gao, PV Interneurons: Critical Regulators of E/I Balance for Prefrontal Cortex-Dependent Behavior and Psychiatric Disorders. Frontiers in neural circuits 12, 37 (2018).

95. C. Minami, T. Shimizu, A. Mitani, Neural activity in the prelimbic and infralimbic cortices of freely moving rats during social interaction: Effect of isolation rearing. PLoS One 12, e0176740 (2017).

96. D. C. Rojas, L. B. Wilson, gamma-band abnormalities as markers of autism spectrum disorders. Biomark Med 8, 353–368 (2014).

97. M. L. Phillips, H. A. Robinson, L. Pozzo-Miller, Ventral hippocampal projections to the medial prefrontal cortex regulate social memory. Elife 8 (2019).

98. R. E. Accordino, C. Kidd, L. C. Politte, C. A. Henry, C. J. McDougle, Psychopharmacological interventions in autism spectrum disorder. Expert Opin Pharmacother 17, 937–952 (2016).

99. D. R. Hampson, S. Gholizadeh, L. K. Pacey, Pathways to drug development for autism spectrum disorders. Clin Pharmacol Ther 91, 189–200 (2012).

100. F. Zhang, C. J. Phiel, L. Spece, N. Gurvich, P. S. Klein, Inhibitory phosphorylation of glycogen synthase kinase-3 (GSK-3) in response to lithium. Evidence for autoregulation of GSK-3. The Journal of biological chemistry 278, 33067–33077 (2003).

101. W. J. Ryves, A. J. Harwood, Lithium inhibits glycogen synthase kinase-3 by competition for magnesium. Biochem Biophys Res Commun 280, 720–725 (2001).

102. W. T. O’Brien, P. S. Klein, Validating GSK3 as an in vivo target of lithium action. Biochemical Society transactions 37, 1133–1138 (2009).

103. D. M. Chuang, H. K. Manji, In search of the Holy Grail for the treatment of neurodegenerative disorders: has a simple cation been overlooked? Biological psychiatry 62, 4–6 (2007).

104. C. Xu, N. G. Kim, B. M. Gumbiner, Regulation of protein stability by GSK3 mediated phosphorylation. Cell cycle 8, 4032–4039 (2009).

105. M. Siegel et al., Preliminary investigation of lithium for mood disorder symptoms in children and adolescents with autism spectrum disorder. J Child Adolesc Psychopharmacol 24, 399–402 (2014).

106. E. M. Anderson et al., Systematic analysis of CRISPR-Cas9 mismatch tolerance reveals low levels of off-target activity. J Biotechnol 211, 56–65 (2015).

107. M. F. Sathler et al., Phosphorylation of the AMPA receptor subunit GluA1 regulates clathrin-mediated receptor internalization. Journal of cell science 134 (2021).

108. K. Sztukowski et al., HIV induces synaptic hyperexcitation via cGMP-dependent protein kinase II activation in the FIV infection model. PLoS biology 16, e2005315 (2018).

109. M. F. Sathler et al., HIV and FIV glycoproteins increase cellular tau pathology via cGMP-dependent kinase II activation. Journal of cell science 135 (2022).

110. S. Kim, A. N. Lapham, C. G. Freedman, T. L. Reed, W. K. Schmidt, Yeast as a tractable genetic system for functional studies of the insulin-degrading enzyme. The Journal of biological chemistry 280, 27481–27490 (2005).

111. S. Kim et al., Brain region-specific effects of cGMP-dependent kinase II knockout on AMPA receptor trafficking and animal behavior. Learning & memory 23, 435–441 (2016).

112. S. Kim et al., Evidence that the rab5 effector APPL1 mediates APP-betaCTF-induced dysfunction of endosomes in Down syndrome and Alzheimer’s disease. Molecular psychiatry 10.1038/mp.2015.97 (2015).

113. S. Kim, J. Shou, S. Abera, E. B. Ziff, Sucrose withdrawal induces depression and anxiety-like behavior by Kir2.1 upregulation in the nucleus accumbens. Neuropharmacology 130, 10–17 (2018).

114. S. Kim et al., Network compensation of cyclic GMP-dependent protein kinase II knockout in the hippocampus by Ca2+-permeable AMPA receptors. Proc Natl Acad Sci U S A 112, 3122–3127 (2015).

115. S. Kim, C. J. Violette, E. B. Ziff, Reduction of increased calcineurin activity rescues impaired homeostatic synaptic plasticity in presenilin 1 M146V mutant. Neurobiol Aging 36, 3239–3246 (2015).

116. S. Kim, E. B. Ziff, Calcineurin mediates synaptic scaling via synaptic trafficking of Ca2+-permeable AMPA receptors. PLoS biology 12, e1001900 (2014).

117. J. P. Roberts, S. A. Stokoe, M. F. Sathler, R. A. Nichols, S. Kim, Selective co-activation of alpha7- and alpha4beta2-nicotinic acetylcholine receptors reverses beta-amyloid-induced synaptic dysfunction. The Journal of biological chemistry 10.1016/j.jbc.2021.100402, 100402 (2021).

118. J. Shou, A. Tran, N. Snyder, E. Bleem, S. Kim, Distinct Roles of GluA2-lacking AMPA Receptor Expression in Dopamine D1 or D2 Receptor Neurons in Animal Behavior. Neuroscience 398, 102–112 (2018).

119. J. L. Sun et al., Co-activation of selective nicotinic acetylcholine receptors is required to reverse beta amyloid-induced Ca(2+) hyperexcitation. Neurobiol Aging 84, 166–177 (2019).

120. T. M. Tran et al., Loss of cGMP-dependent protein kinase II alters ultrasonic vocalizations in mice, a model for speech impairment in human microdeletion 4q21 syndrome. Neuroscience letters 759, 136048 (2021).

121. A. Zaytseva et al., Ketamine’s rapid antidepressant effects are mediated by Ca^2+^-permeable AMPA receptors in the hippocampus. bioRxiv 10.1101/2022.12.05.519102, 2022.2012.2005.519102 (2022).

122. J. Y. Kim et al., Viral transduction of the neonatal brain delivers controllable genetic mosaicism for visualising and manipulating neuronal circuits in vivo. The European journal of neuroscience 37, 1203–1220 (2013).

123. H. Dana et al., High-performance calcium sensors for imaging activity in neuronal populations and microcompartments. Nature methods 16, 649–657 (2019).

124. J. Dimidschstein et al., A viral strategy for targeting and manipulating interneurons across vertebrate species. Nat Neurosci 19, 1743–1749 (2016).

125. S. Kim et al., Neural circuit pathology driven by Shank3 mutation disrupts social behaviors. Cell Rep 39, 110906 (2022).

126. S. Kim, Y. E. Kim, I. H. Kim, Simultaneous analysis of social behaviors and neural responses in mice using round social arena system. STAR Protoc 3, 101722 (2022).

