## Supplementary Tables for "The autism-associated loss of δ-catenin functions disrupts social behaviors"

**Table S1. Three-chamber test summary of WT and G34S mice with regular and lithium chow**

| Regular chow | | | | | | | | | |
| --- | --- | --- | --- | --- | --- | --- | --- | --- | --- |
| Female | WT (n = 11 mice) | | G34S (n = 8 mice) | | Male | WT (n = 11 mice) | | G34S (n = 14 mice) | |
| Sociability | Stranger 1 | Objective | Stranger 1 | Objective | Sociability | Stranger 1 | Objective | Stranger 1 | Objective |
| Total Interaction Time (sec) | 16.445 ± 4.244 | 10.855 ± 2.973 | 11.538 ± 3.387 | 13.513 ± 3.433 | Total Interaction Time (sec) | 16.864 ± 5.169 | 8.655 ± 3.244 | 10.986 ± 4.035 | 10.829 ± 2.691 |
|  | Stranger 1 vs. Objective,  *p* = 0.0042 | | Stranger 1 vs. Objective,  *p* = 0.3888 | |  | Stranger 1 vs. Objective,  *p* < 0.0001 | | Stranger 1 vs. Objective,  *p* = 0.9995 | |
|  | WT Stranger 1 vs. G34S Stranger 1, *p* = 0.0268 | | | |  | WT Stranger 1 vs. G34S Stranger 1, *p* = 0.0023 | | | |
| Social novelty | Stranger 1 | Stranger 2 | Stranger 1 | Stranger 2 | Social novelty | Stranger 1 | Stranger 2 | Stranger 1 | Stranger 2 |
| Total Interaction Time (sec) | 6.182 ± 1.795 | 12.827 ± 4.303 | 6.825 ± 2.712 | 8.425 ± 2.843 | Total Interaction Time (sec) | 7.391 ± 3.428 | 17.573 ± 4.037 | 7.679 ± 9.314 | 9.314 ± 5.332 |
|  | Stranger 1 vs. 2,  *p* < 0.0001 | | Stranger 1 vs. 2,  *p* = 0.7307 | |  | Stranger 1 vs. 2,  *p* < 0.0001 | | Stranger 1 vs. 2,  *p* = 0.7297 | |
|  | WT Stranger 2 vs. G34S Stranger 2, *p* = 0.0212 | | | |  | WT Stranger 2 vs. G34S Stranger 2, *p* < 0.0001 | | | |
| Lithium chow | | | | | | | | | |
| Female | WT (n = 13 mice) | | G34S (n = 9 mice) | | Male | WT (n = 8 mice) | | G34S (n = 12 mice) | |
| Sociability | Stranger 1 | Objective | Stranger 1 | Objective | Sociability | Stranger 1 | Objective | Stranger 1 | Objective |
| Total Interaction Time (sec) | 36.192 ± 14.596 | 15.069 ± 7.317 | 31.800 ± 14.691 | 13.544 ± 4.571 | Total Interaction Time (sec) | 32.850 ± 12.862 | 15.500 ± 9.699 | 28.042 ± 10.979 | 16.667 ± 7.563 |
|  | Stranger 1 vs. Objective,  *p* = 0.0001 | | Stranger 1 vs. Objective,  *p* = 0.0074 | |  | Stranger 1 vs. Objective,  *p* = 0.0089 | | Stranger 1 vs. Objective,  *p* = 0.0468 | |
|  | WT Stranger 1 vs. G34S Stranger 1, *p* = 0.8061 | | | |  | WT Stranger 1 vs. G34S Stranger 1, *p* = 0.7336 | | | |
| Social novelty | Stranger 1 | Stranger 2 | Stranger 1 | Stranger 2 | Social novelty | Stranger 1 | Stranger 2 | Stranger 1 | Stranger 2 |
| Total Interaction Time (sec) | 8.746 ± 3.242 | 33.154 ± 19.031 | 10.433 ± 7.117 | 35.989 ± 13.988 | Total Interaction Time (sec) | 7.363 ± 3.864 | 18.800 ± 12.404 | 8.450 ± 4.345 | 17.258 ± 5.579 |
|  | Stranger 1 vs. 2,  *p* < 0.0001 | | Stranger 1 vs. 2,  *p* = 0.0006 | |  | Stranger 1 vs. 2,  *p* = 0.0117 | | Stranger 1 vs. 2,  *p* = 0.0182 | |
|  | WT Stranger 2 vs. G34S Stranger 2, *p* = 0.9550 | | | |  | WT Stranger 2 vs. G34S Stranger 2, *p* = 0.9615 | | | |

**Table S2. The discrimination index of WT and G34S mice with regular and lithium chow**

| Female | Regular chow | | Lithium chow | | Male | Regular chow | | Lithium chow | |
| --- | --- | --- | --- | --- | --- | --- | --- | --- | --- |
|  | WT | G34S | WT | G34S |  | WT | G34S | WT | G34S |
| Sociability Discrimination index | 0.203 ± 0.126 | -0.087 ± 0.128 | 0.401 ± 0.150 | 0.383 ± 0.081 | Sociability Discrimination index | 0.323 ± 0.201 | -0.014 ± 0.226 | 0.389 ± 0.131 | 0.267 ± 0.147 |
|  | WT vs. G34S,  *p* = 0.0001 | | WT vs. G34S,  *p* = 0.9868 | |  | WT vs. G34S,  *p* = 0.0003 | | WT vs. G34S,  *p* = 0.4847 | |
|  | G34S regular chow vs. G34S lithium, *p* < 0.0001 | | | |  | G34S regular chow vs. G34S lithium, *p* =0.0024 | | | |
| Social Novelty Discrimination index | 0.341 ± 0.091 | 0.107 ± 0.134 | 0.532 ± 0.166 | 0.548 ± 0.227 | Social Novelty Discrimination index | 0.423 ± 0.208 | 0.015 ± 0.290 | 0.424 ± 0.262 | 0.379 ± 0.171 |
|  | WT vs. G34S,  *p* = 0.0168 | | WT vs. G34S,  *p* = 0.9954 | |  | WT vs. G34S,  *p* = 0.0115 | | WT vs. G34S,  *p* = 0.0115 | |
|  | G34S regular chow vs. G34S lithium, *p* < 0.0001 | | | |  | G34S regular chow vs. G34S lithium, *p* =0.0372 | | | |

**Table S3. Summary of the buried food and open field tests**

|  | Buried food test | Open field test | | |
| --- | --- | --- | --- | --- |
| Female | The latency (Seconds) | Total distance travelled (m) | Total time spent outside (Seconds) | Total time spent inside (Seconds) |
| WT | 76.000 ± 22.443 | 52.378 ± 6.751 | 1045.150 ± 54.380 | 154.850 ± 54.380 |
| G34S | 83.727 ± 65.521 | 48.454 ± 13.409 | 1062.338 ± 70.804 | 154.892 ± 73.942 |
|  | *p* = 0.7694 | *p* = 0.4537 | *p* = 0.5645 | *p* = 0.9988 |
| Male | The latency (Seconds) | Total distance travelled (m) | Total time spent outside (Seconds) | Total time spent inside (Seconds) |
| WT | 86.200 ± 63.554 | 53.532 ± 12.868 | 1033.308 ± 63.382 | 166.692 ± 63.382 |
| G34S | 90.333 ± 60.593 | 51.742 ± 7.954 | 1037.368 ± 69.715 | 162.632 ± 69.715 |
|  | *p* = 0.8777 | *p* = 0.6346 | *p* = 0.8713 | *p* = 0.8623 |
|  | Buried food test | Open field test | | |
| Female | The latency (Seconds) | Total distance travelled (m) | Total time spent outside (Seconds) | Total time spent inside (Seconds) |
| WT | 180.250 ± 175.360 | 65.701 ± 16.926 | 1028.536 ± 57.074 | 171.464 ± 57.074 |
| KO | 178.833 ± 135.632 | 65.285 ± 18.005 | 1005.225 ± 42.884 | 194.775 ± 42.884 |
|  | *p* = 0.9864 | *p* = 0.9573 | *p* = 0.3288 | *p* = 0.3288 |
| Male | The latency (Seconds) | Total distance travelled (m) | Total time spent outside (Seconds) | Total time spent inside (Seconds) |
| WT | 185.542 ± 130.374 | 51.071 ± 20.755 | 1054.510 ± 75.972 | 145.490 ± 75.972 |
| KO | 165.200 ± 141.551 | 59.102 ± 18.124 | 1057.845 ± 47.091 | 142.155 ± 47.091 |
|  | *p* = 0.6885 | *p* = 0.2909 | *p* = 0.8961 | *p* = 0.8961 |

**Table S4. Three-chamber test summary of WT and KO mice.**

| Regular chow | | | | | | | | | |
| --- | --- | --- | --- | --- | --- | --- | --- | --- | --- |
| Female | WT (n = 16 mice) | | KO (n = 14 mice) | | Male | WT (n = 20 mice) | | KO (n = 15 mice) | |
| Sociability | Stranger 1 | Objective | Stranger 1 | Objective | Sociability | Stranger 1 | Objective | Stranger 1 | Objective |
| Total Interaction Time (sec) | 16.981 ± 8.908 | 8.731 ± 3.853 | 9.393 ± 5.173 | 9.829 ± 2.496 | Total Interaction Time (sec) | 19.225 ± 12.543 | 8.290 ± 4.968 | 11.053 ± 6.531 | 9.840 ± 3.052 |
|  | Stranger 1 vs. Objective,  *p* = 0.0008 | | Stranger 1 vs. Objective,  *p* = 0.9971 | |  | Stranger 1 vs. Objective,  *p* = 0.0003 | | Stranger 1 vs. Objective,  *p* = 0.9753 | |
|  | WT Stranger 1 vs. KO Stranger 1, *p* = 0.0035 | | | |  | WT Stranger 1 vs. KO Stranger 1, *p* = 0.0192 | | | |
| Social novelty | Stranger 1 | Stranger 2 | Stranger 1 | Stranger 2 | Social novelty | Stranger 1 | Stranger 2 | Stranger 1 | Stranger 2 |
| Total Interaction Time (sec) | 3.488 ± 3.466 | 16.994 ± 10.683 | 4.293 ± 4.296 | 7.221 ± 5.148 | Total Interaction Time (sec) | 3.670 ± 2.497 | 20.880 ± 11.455 | 6.713 ± 5.899 | 9.380 ± 4.679 |
|  | Stranger 1 vs. 2,  *p* < 0.0001 | | Stranger 1 vs. 2,  *p* = 0.6511 | |  | Stranger 1 vs. 2,  *p* < 0.0001 | | Stranger 1 vs. 2,  *p* = 0.7403 | |
|  | WT Stranger 2 vs. KO Stranger 2, *p* = 0.0010 | | | |  | WT Stranger 2 vs. KO Stranger 2, *p* < 0.0001 | | | |

**Table S5. The discrimination index of WT and KO mice**

| Regular chow | | | | | |
| --- | --- | --- | --- | --- | --- |
| Female | WT | KO | Male | WT | KO |
| Sociability Discrimination index | 0.307 ± 0.150 | -0.081 ± 0.373 | Sociability Discrimination index | 0.360 ± 0.195 | -0.011 ± 0.408 |
|  | *p* = 0.0007 | |  | *p* = 0.0010 | |
| Social Novelty Discrimination index | 0.676 ± 0.222 | 0.322 ± 0.405 | Social Novelty Discrimination index | 0.657 ± 0.232 | 0.255 ± 0.412 |
|  | *p* = 0.0053 | |  | *p* = 0.0009 | |

**Table S6. Three-chamber test summary of G34S and KO mice with lithium chow.**

| Lithium chow | | | | | | | | | |
| --- | --- | --- | --- | --- | --- | --- | --- | --- | --- |
| Female | G34S (n = 9 mice) | | KO (n = 7 mice) | | Male | G34S (n = 12 mice) | | KO (n = 7 mice) | |
| Sociability | Stranger 1 | Objective | Stranger 1 | Objective | Sociability | Stranger 1 | Objective | Stranger 1 | Objective |
| Total Interaction Time (sec) | 31.800 ± 14.691 | 31.800 ± 14.691 | 13.200 ± 5.026 | 13.571 ± 4.549 | Total Interaction Time (sec) | 28.042 ± 10.979 | 16.667 ± 7.563 | 17.771 ± 2.906 | 13.457 ± 3.363 |
|  | Stranger 1 vs. Objective,  *p* = 0.0008 | | Stranger 1 vs. Objective,  *p* = 0.9998 | |  | Stranger 1 vs. Objective,  *p* = 0.0058 | | Stranger 1 vs. Objective,  *p* = 0.7312 | |
|  | WT Stranger 1 vs. G34S Stranger 1, *p* = 0.0014 | | | |  | WT Stranger 1 vs. G34S Stranger 1, *p* = 0.0431 | | | |
| Social novelty | Stranger 1 | Stranger 2 | Stranger 1 | Stranger 2 | Social novelty | Stranger 1 | Stranger 2 | Stranger 1 | Stranger 2 |
| Total Interaction Time (sec) | 10.433 ± 7.117 | 35.989 ± 13.988 | 15.371 ± 6.336 | 20.500 ± 6.562 | Total Interaction Time (sec) | 8.958 ± 4.248 | 16.650 ± 5.767 | 14.443 ± 6.724 | 17.300 ± 3.818 |
|  | Stranger 1 vs. 2,  *p* < 0.0001 | | Stranger 1 vs. 2,  *p* = 0.7383 | |  | Stranger 1 vs. 2,  *p* = 0.0011 | | Stranger 1 vs. 2,  *p* = 0.7310 | |
|  | WT Stranger 2 vs. G34S Stranger 2, *p* = 0.0142 | | | |  | WT Stranger 2 vs. G34S Stranger 2, *p* > 0.9999 | | | |

**Table S7. The discrimination index of G34S and KO mice with lithium chow**

| Lithium chow | | | | | |
| --- | --- | --- | --- | --- | --- |
| Female | G34S | KO | Male | G34S | KO |
| Sociability Discrimination index | 0.382 ± 0.081 | -0.017 ± 0.130 | Sociability Discrimination index | 0.267 ± 0.147 | 0.111 ± 0.102 |
|  | *p* < 0.0001 | |  | *p* = 0.0247 | |
| Social Novelty Discrimination index | 0.549 ± 0.228 | 0.147 ± 0.304 | Social Novelty Discrimination index | 0.379 ± 0.171 | 0.116 ± 0.168 |
|  | *p* = 0.0090 | |  | *p* = 0.0046 | |
